# The Consciousness Theories Studies (ConTraSt) database: analyzing and comparing empirical studies of consciousness theories

**DOI:** 10.1101/2021.06.10.447863

**Authors:** Itay Yaron, Lucia Melloni, Michael Pitts, Liad Mudrik

**Author notes:** Corresponding Author: Itay Yaron, The Sagol school of neuroscience and the school of Psychological sciences, Tel Aviv University, Rama Aviv, P.O.B 39040, Tel Aviv, ISRAEL.

## Abstract

Understanding how consciousness arises from neural activity remains one of the biggest challenges for neuroscience. Numerous theories have been proposed in recent years, each gaining independent empirical support. Currently, there is no comprehensive, quantitative and theory-neutral overview of the field that enables an evaluation of how theoretical frameworks interact with empirical research. We provide a bird’s eye view on studies that interpreted their findings in light of at least one of four leading neuroscientific theories of consciousness (N=412 experiments), asking how methodological choices of the researchers might affect the final conclusions. We found that supporting a specific theory can be predicted solely from methodological choices, irrespective of findings. Furthermore, most studies interpret their findings post-hoc, rather than a-priori testing critical predictions of the theories. Our results highlight challenges for the field and provide researchers with a unique, open-access website to further analyze trends in the neuroscience of consciousness.

## Main

Within about three decades, the scientific study of consciousness has transitioned from an emerging field, trying to establish its legitimacy, to a flourishing source of empirical studies and theoretical accounts. Concomitant with the empirical quest towards finding the neural correlates of consciousness^1,2^, the field has seen a steep growth in the number of scientific theories, each offering a different explanation of the neural basis of consciousness^3–6^. Accordingly, the field has not yet converged around an accepted account, and disagreements abound even with respect to the neural correlates themselves ^7–10^. Existing reviews of the state-of-the-art of the field are typically written by proponents of the different theories^11–13^, with findings described predominantly through the lens of a given theoretical framework, yielding dramatically different pictures.

Converging onto a unified account is even harder given the wide range of experimental paradigms used to study consciousness, which might systematically differ in studies supporting the different theories. Among other methods, common experimental procedures include Masking ^14^, Bistable perception ^15,16^, Inattentional Blindness^17^, Change Blindness ^18^, Stimulus Degradation ^19^, Sleep ^20^, Anesthesia ^21^, Direct Stimulation (e.g., by TMS^22^, intracranial stimulation ^23^, etc.). Similarly, different measures of consciousness exist (e.g., report vs. no-report paradigms^24^). These different procedures probe somewhat different processes^3^, sometimes yielding different results^25–27^. Thus, if there is a systematic bias in methodological choices, this could explain how conflicting theories appear to be supported by empirical data, despite the proclaimed goal to test and account for the very same phenomenon^3^. Critically, the choice of paradigm or analysis approach might affect the conclusions one draws. For example, in the recent debate around the role of prefrontal cortex in consciousness, both sides suggested the conclusions of their opponents were based on problematic methodological choices^9,28^ (for other debates, see ^24,29^ or ^30,31^).

Here, we present an unbiased, theory-neutral, quantitative and systematic review of empirical findings around leading theories of consciousness, providing a bird’s eye view of the field and looking for potential biases in interpreting empirical findings. We focus on four theories that have evoked substantial empirical and theoretical interest ^3,4,8^: Global Neuronal Workspace^12,32^ (GNW), Higher-order thought^13,33^ (HOT), Integrated Information Theory^34,35^ (IIT), and Recurrent Processing Theory^36,37^ (RPT) (listed in alphabetical order). The theories differ in their core principles, suggested mechanisms, and predictions they make about neural activity associated with consciousness^3^. In a nutshell (for a more detailed description see Supplementary Box A), GNW claims that conscious processing emerges when information is globally broadcasted by a frontoparietal network^12,38^, while HOT ascribes consciousness to higher-order representations in dorsolateral prefrontal cortex, that accompany first-order representations elsewhere^13^. IIT, conversely, equates consciousness with a maximum of irreducible intrinsic cause–effect power, as determined from the intrinsic perspective of the system, and claims that a local maximum of such power likely resides in a posterior cortical “hot zone”^34,39,40^. Finally, RPT asserts that horizontal connections and recurrent loops between lower and higher-level brain areas, involving plastic changes mediated by NMDA-dependent feedback activations, underlie conscious processing^11,36,37^.

As a first step towards building a robust dataset of relevant empirical findings, we collected papers that interpreted their results as supporting or challenging GNW, HOT, IIT, or RPT. All collected papers were classified according to various parameters of interest, including, among others, the experimental paradigms, stimuli, neuroscientific techniques, empirical findings, and theoretical interpretations (see Supplementary Table S1 for the full parameter list). Using these parameters, we provide a descriptive overview of the field, in addition to a data-driven statistical analysis, aimed to uncover trends, biases, blind spots, and limitations regarding how the theories and the empirical studies interact.

To reveal such biases, we conducted an epistemic, meta-experimental examination of the field, asking how findings are interpreted and to what extent supporting a theory depends on methodological decisions made by researchers. Rather than conducting a standard meta-analysis aimed at identifying consistent effects and providing consolidated estimates of such effects, we focused on the way these findings are collected and interpreted. Thus, we took the reported findings at face value, with no attempt to re-interpret or test the statistical reliability of the findings, or to critically assess the strength of the experimental design. This allowed us to identify what claims are being made and what types of findings and paradigms these claims are based on. Sticking to the original interpretations of the authors also allowed us to keep this review objective and uncontaminated by subjective judgments of methodological quality or the correct interpretation of the findings. An additional objective was to provide the scientific community with an interactive, open-access online tool to gauge the state of the field and the status of the leading theories. This rich database encompasses 412 experiments reported in 365 studies published between April 2001 and October 2019.

## Results

The database was created following predefined criteria (see methods) in line with the PRISMA 2009 guidelines^41^ (Figure 1). Our search strategy looked for papers between the years 2001 and 2019 (up until the search date of October 22), relying on (a) *Topic search*, including papers in which the name of each of the four theories appears as part of the topic / abstract / keywords; and (b) *Citation search*, where we first identified three key papers for each theory (Supplementary Table S1), and then, collected all empirical papers citing one or more of these twelve key papers.

**Figure 1.**
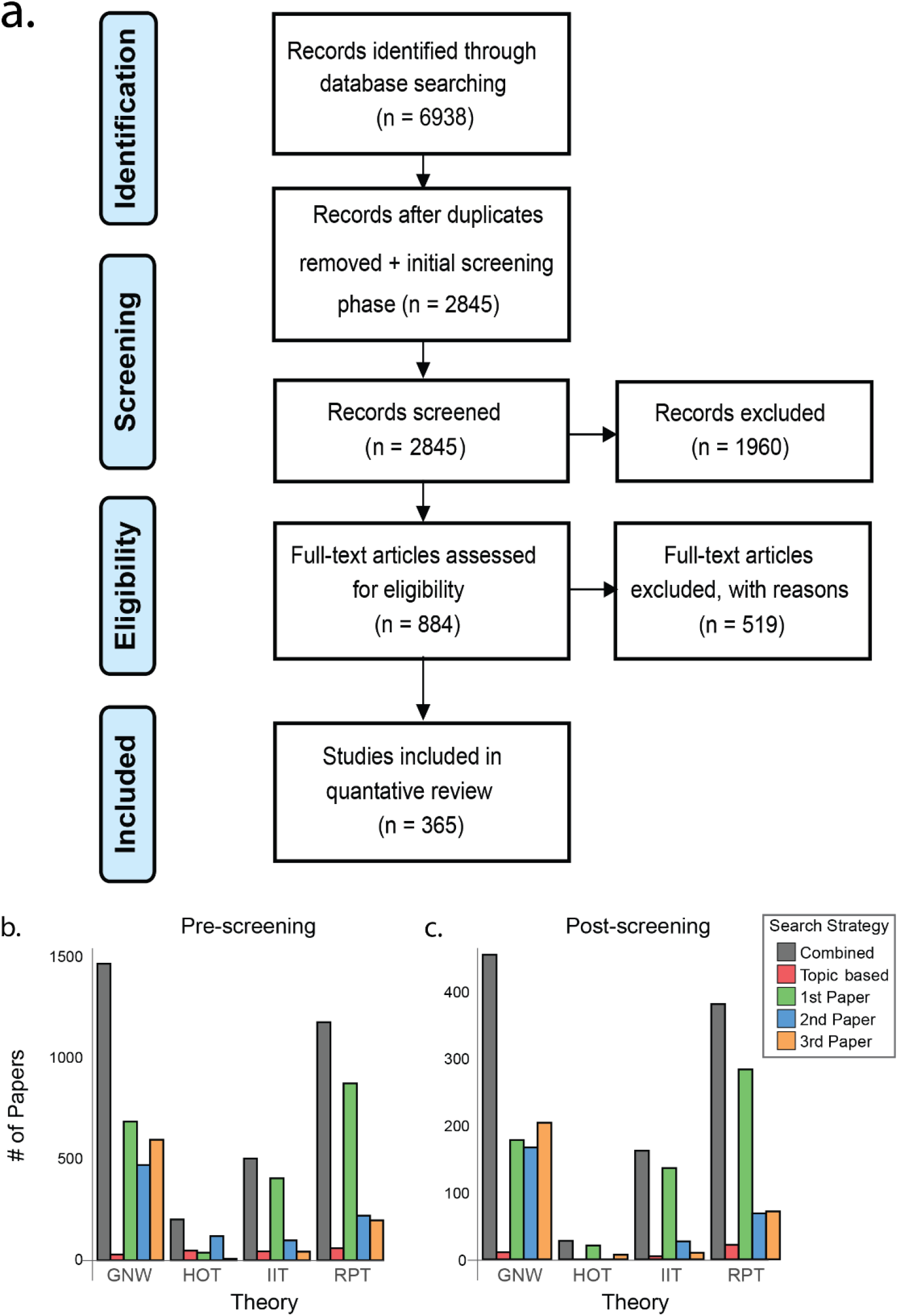
Panel a: Flow diagram outlining the process of article selection^41^. Panel b: Cumulative distribution of the papers found for each of the theories pre-screening, according to the different search strategies. Panel c: Cumulative distribution of the final papers post-screening. For panels b & c, each bar depicts the number of unique papers in that category. Gray bars represent all papers found for each theory across the different search strategies. Red bars denote papers that mention the theory in their title, abstract, or keywords. Green, blue, and orange bars denote papers citing the first, second, and third key papers for each theory (see Supplementary Table S1 for the key theory papers).

Out of 6938 records identified in the initial database searching, 6054 were filtered out due to predefined constraints (for a full list, see again methods). The remaining 884 unique papers were assessed for eligibility by a close inspection of their full-text, removing papers that: (1) did not directly relate to consciousness (n = 232; see Supplementary Figure S1 and Supplementary Table S2 for the distribution of excluded papers into main fields of research); (2) did not interpret their results in light of an NCC prediction of one or more of the theories (n = 190); (3) Reviews (n = 89); (4) Meta-analyses (n = 5); and (5) Behavioral studies (n = 3). This process yielded the final database, which included 365 papers, reporting 412 experiments, all of which were classified according to our predefined parameters of interest (see Supplementary Table S3).

### Division of experiments with respect to the theories

The most prominent finding was the non-uniform distribution of experiments mentioning each of the four theories (Figure 2a; note that this might be explained – at least in part – by the different ‘age’ of the theories, see again Supplementary Table S1). Namely, in this database, GNW is most widely discussed (N=224), followed by RPT (N=140), IIT (N= 101), and HOT (N=12; note that for HOT, our initial search yielded a much larger number of papers (N= 532), but most of them did not survive the screening process as they were either not empirical, or did not include neuroscientific results; see Methods). Notably, the distribution is also highly skewed with respect to confirmatory (experiments supporting the theories) vs. disconfirmatory experiments (experiments challenging the theories), the latter constituting only 15% of all experiments. As evident in Figure 2a, this skewed pattern was observed for GNW (*χ*^2^ (1) = 90.02, *p* < .0001), RPT (*χ*^2^ (1) = 77.26, *p* < .0001) and IIT(*χ*^2^ (1) = 78.43, *p* < .0001), but not for HOT (*χ*^2^ (1) = 0.33, *p* = .60; note again the low representation of HOT in our database). An inspection of the distribution over time suggests that all theories have been increasingly studied (Figure 2b) and have gained support (Figure 2c) through the years. Interestingly, there does seem to be some rise in challenging experiments in the last decade (especially for GNW), that might reflect a gradual maturation of the field (Figure 2d). However, visually inspecting the trends over time suggests that the increasing support of each theory is unaffected by the changing support of other theories, demonstrating a parallel progression of leading theories. That is, the number of experiments referring to the theories keeps growing, as opposed to a replacement model, where the growing success of one theory leads to a gradual reduction of support for the other (e.g., ^42^, when one theory reductively replaces the other), which would translate into a plateau-like trend, where no new studies are added.

**Figure 2.**
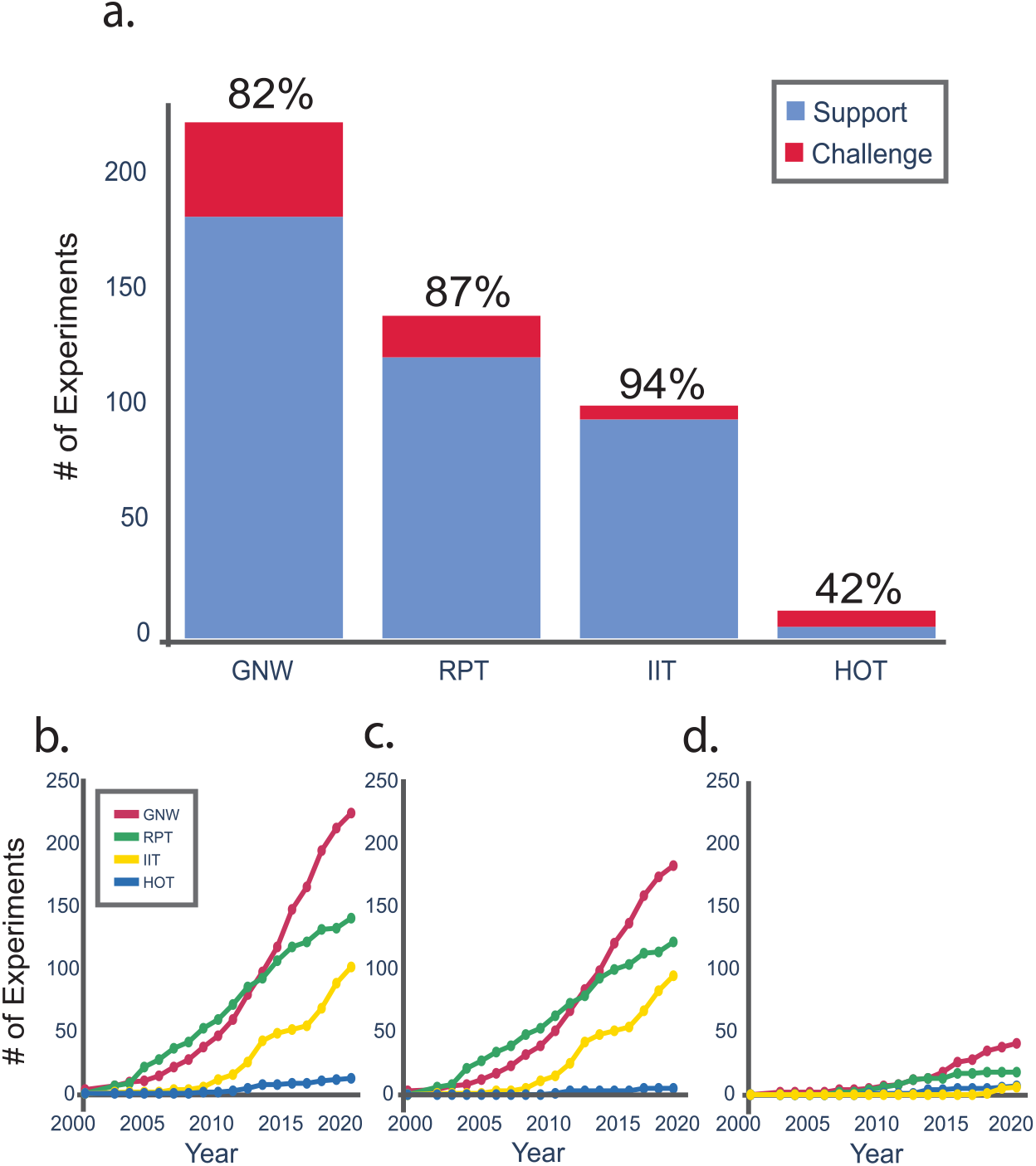
Panel a: Distribution of experiments across theories. Each bar represents the number of experiments interpreted as supporting the theory (blue) or challenging it (red). The label above each bar indicates the proportion of papers supporting each theory. Panel b: Cumulative distribution over time of experiments studying/discussing each of the theories (either challenging or supporting them). Panel c: Cumulative distribution over time of the experiments *supporting* the theories. Panel d: Cumulative distribution over time of the experiments *challenging* the theories.

Notably, only about a third (35%) of the experiments were presented as explicitly testing theory predictions, as opposed to 41% of the experiments that interpreted their findings in light of the theories post-hoc, in the discussion section (the remaining 24% generally mentioned a theory in the introduction, without formulating clear hypotheses about it and without interpreting the evidence post-hoc). Importantly, only 7% tested predictions of more than one theory, trying to pit them against each other. Another important observation was that theory-driven experiments, post-hoc-interpreted experiments, and experiments that only mentioned the theories in the introduction differed in their likelihood to challenge a theory (note that in this analysis, papers were classified as challenging a theory if at least one theory was challenged, even if another theory was supported, while the rest were classified as ‘not challenging’) (*χ*^2^ (2) = 68.79, *p* < .0001; Figure 3). Theory-driven experiments challenged the theories more frequently than expected under the null hypothesis (*Z* = 6.14, *p* < .001) and the post-hoc-interpreting/theory-mentioning papers challenged them less frequently than expected (*Z* = − 4.07, *p* < .001)and marginally, *Z* = −2.05, *p* = .06, respectively).

**Figure 3.**
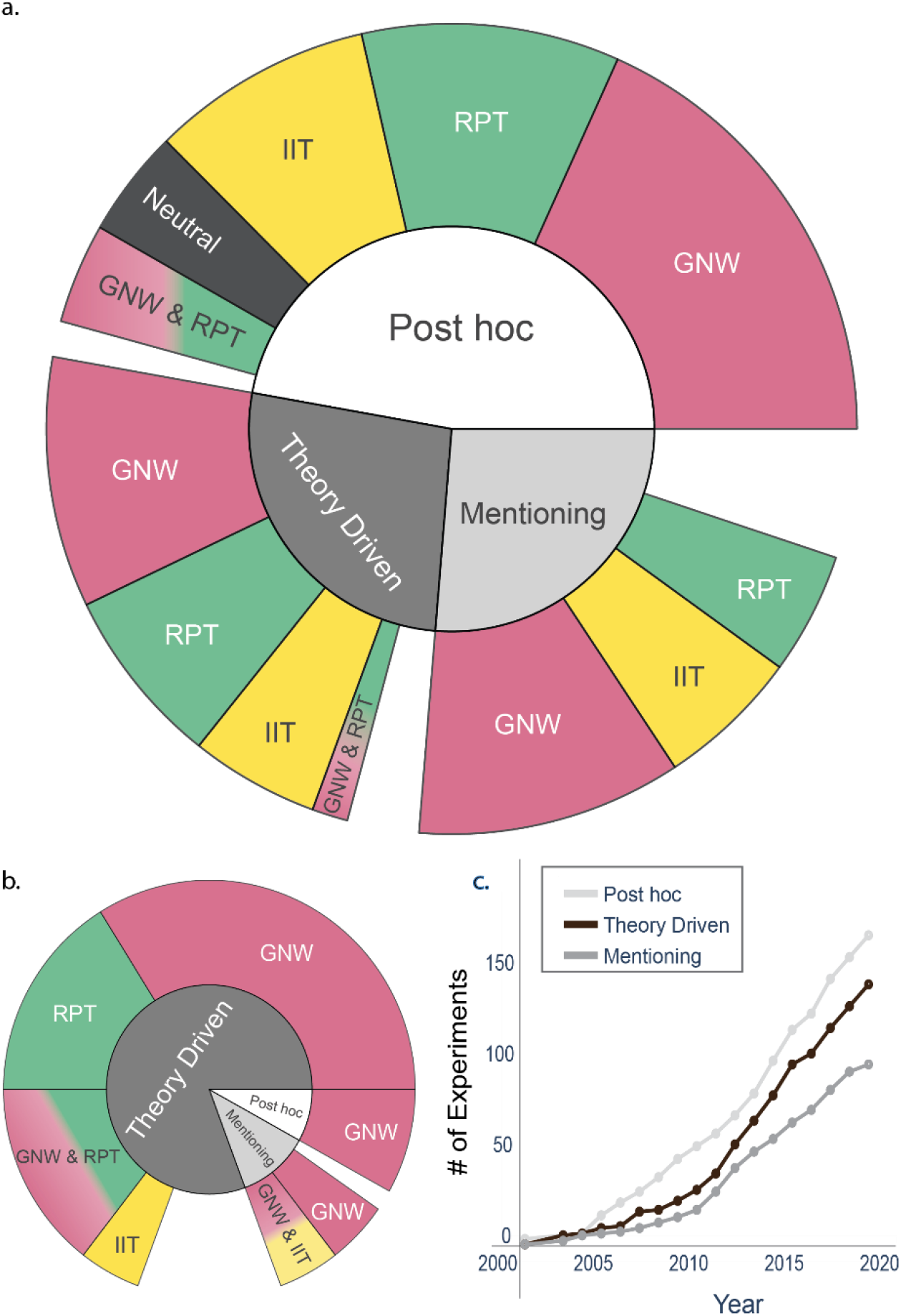
Panel a: Distribution of experiments that did not challenge any theory (N=350) divided into “theory-driven” (i.e., explicitly testing at least one prediction of at least one theory; dark gray in the inner circle), “mentioning” at least one theory in their introduction (light gray), or “post-hoc” interpreting their results in light of at least one theory, without referring to it in the introduction (white). The outer circle describes the distribution of experiments to specific theories. Panel b: Similar to Panel a, but for experiments interpreted as challenging at least one theory (N=62). Slices with less than five/three experiments (e.g., studies challenging HOT) were not included in Panel a and Panel b, respectively (hence the empty spaces in the outer circles). Panel c: a cumulative distribution of all experiments that are either theory-driven (black), mention the theories (dark gray) or interpret the findings post hoc (light gray) over time.

### Prediction of supported theory based on methodological parameters

To assess the potential influence of methodological choices on the probability of a study supporting/challenging a specific theory, a random forest classifier was used. Specifically, we trained a classifier to predict whether each experiment will support GNW, IIT, RPT or any combination of these theories (HOT was not included due to insufficient number of experiments). The classifier used all methodological parameters in our database, excluding parameters showing multicollinearity ^43^ (for the list of included parameters, see Supplementary Table S5; for the results of the full model without exclusion, see Supplementary Figure S2a-b and Supplementary Table S6). A leave-one-out strategy was used to measure the accuracy of the classifier, which was 80.34%, *t*(411)= 10.99, *p* < 0.001, with a chance level at 67.64% (chance level was determined based on the experiments’ marginal distributions to theories (see Methods), so the accuracy of the tested model is compared with a model which solely relies on the frequency of the outcome in our database, without taking into account any of the parameters (Figure 4a). This accordingly corrects for the different number of experiments for each theory). Our sensitivity analysis (see Methods) validated these results, showing stable and high classification accuracies across different random states of the classifier (M = 79.98%, SD = .42%, range: 78.64%-81.31%). Signal Detection Theory (SDT) analysis^44^ revealed that the classifier performed well for all three theories, with an area under the curve (AUC) of 0.7, 0.84, and 0.76 for GNW, IIT, and RPT respectively (Figure 4b). The individual importance of the features on which the classifier was trained was calculated using a permutation importance method (see Methods). Parameters whose average importance was higher than 95% of the distribution of importance scores of a random parameter were (in order of importance): (1) studying state vs. content consciousness; (2) using report vs. no-report paradigms; (3); using connectivity measures and (4) using subjective measures of consciousness (Supplementary Table S5, Supplementary Figure S3). Below we elaborate on each of these factors, and include HOT for all descriptive statistics.

**Figure 4.**
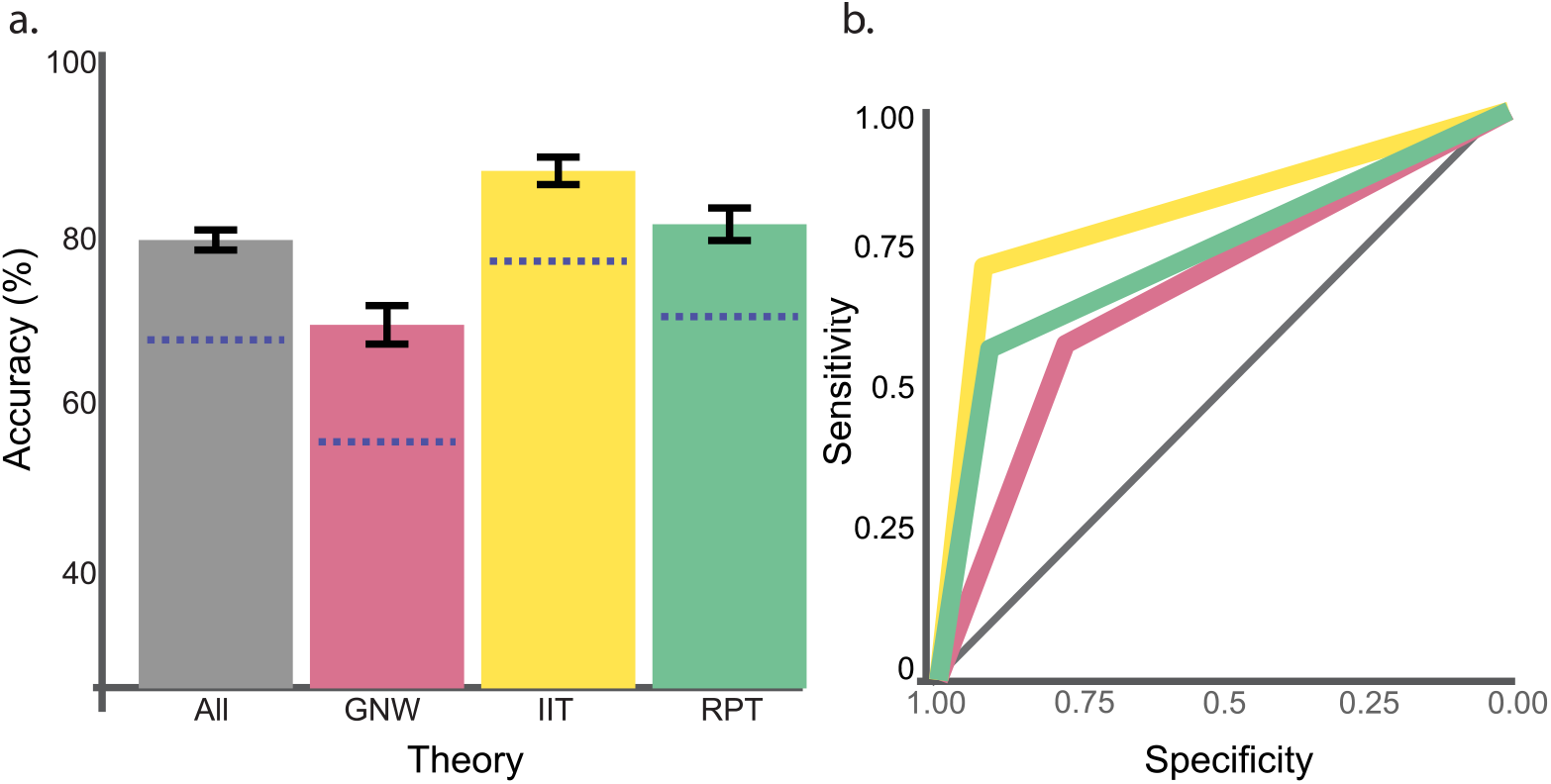
Analysis of the classifier’s performance in predicting support for each theory based on methodological choices. Panel a: classification accuracy for the three theories combined (gray), and for each theory: GNW (red), IIT (yellow), and RPT (green). Panel b: ROC curves for classification performance for each of the theories. Blue dashed lines in Panel a indicate chance level performance for each of the conditions, assessed based on the marginal distributions of the experiments to theories, and gray diagonal line in Panel b indicates the performance of a random classifier (see Methods).

Out of the factors identified as important in classifying the supported theory, the most striking one is content vs. state studies (Figure 5a); experiments supporting IIT are mostly focused on state consciousness (79%), while the exact opposite pattern appears for GNW (73% focusing on content consciousness) and is even more extreme for HOT and RPT (100% and 97% focus on content consciousness). Along the same lines, report paradigms are more prevalent for GNW, RPT, and HOT (58%, 80%, 100%, respectively; note that these numbers are calculated across state and content studies), while IIT gains more support from no-report paradigms (85%) (Figure 5b). Notably, this pattern is largely driven by experiments studying state consciousness, yet it is also observed – to a lesser degree – in studies of content consciousness (Figure 5c). Connectivity dependent measures are more frequently used in experiments supporting IIT (64%) and GNW (37%), compared with RPT (13%) (Figure 5d). Lastly, for the factor ‘measure of consciousness’, objective measures of consciousness (i.e., judgements that can either be correct or incorrect about the critical stimulus^45^) were generally more prevalent than subjective ones (i.e., reporting the degree to which the stimulus was consciously experienced^46,47^; 50% vs. 44% overall, and 69% vs. 61% within content consciousness, respectively). This trend was mostly found for RPT and GNW (73% objective vs. 51% subjective for RPT, and 51% vs. 44% for GNW, and within experiments studying content (as opposed to state) consciousness 75% vs. 53%, 69% vs. 59%), which was not the case for IIT (11% vs. 16%, and within content consciousness 57% vs. 71%) (Figure 5e).

**Figure 5.**
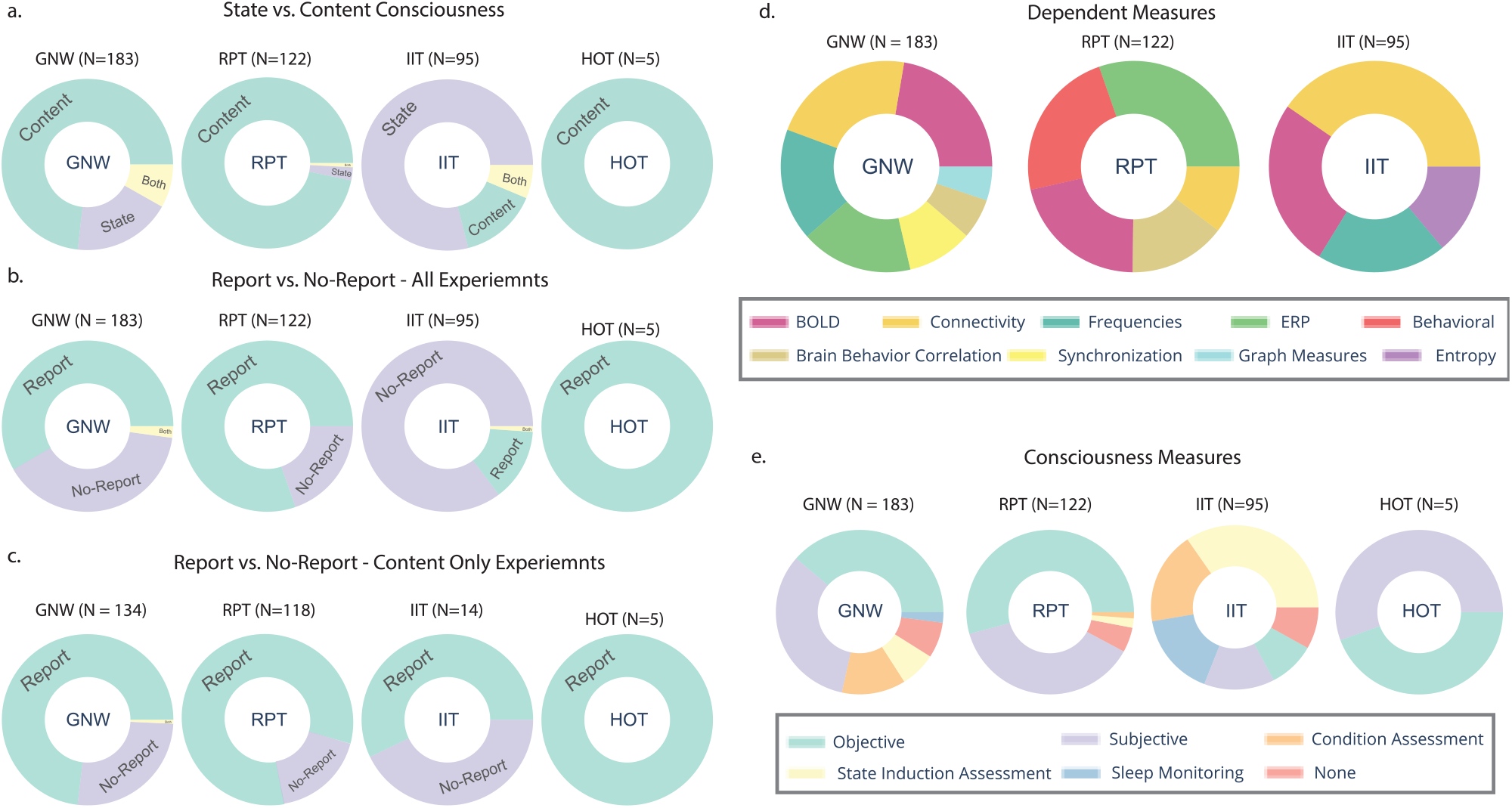
Distribution of experiments supporting a certain theory according to specific methodological parameters: a: Consciousness type; b: Report vs. no-report paradigms; c: similar to Panel b, including only experiments that studied ‘content’ consciousness. d: Type of dependent measures. e. Measures of consciousness. Slices with less than 15 experiments were not included in panel d. Abbreviations: BOLD (Blood-oxygen-level-dependent), ERP (event-related potentials), BBC (brain-behavior correlation). Note that some experiments used more than one consciousness measure or reported more than one type of dependent measure, so they could appear in more than one slice in panels d and e.

### Experimental procedures and techniques

Beyond identifying potential methodological biases with respect to the theories, our database allows for inspection of trends in the field, with respect to preferred paradigms, techniques, and common practices. The most frequently used experimental paradigms (see Supplementary Figure S4a) across all experiments is stimulus degradation, where the strength of the stimulus is reduced (e.g., using contrast^48^, noise^49^ or coherence manipulation^50^) (22%) followed by masking (18%) and direct stimulation (e.g., TMS, tDCS, tACS, intracranial stimulation; 17%). When dividing the experiments to studies focusing on content (Supplementary Figure S4b) or state (Supplementary Figure S4c) consciousness, two very different distributions emerge, as could be expected. A single exception for this pattern is the use of direct stimulation, which was prevalent in both distributions. Given the wide heterogeneity of methods in the field, and especially between state and content studies, we further conducted a post-hoc exploratory analysis, where we asked whether *within each type* (i.e., content vs. state consciousness), the choice of paradigm could predict support for different theories. We used the same random forest method on experiments investigating content or state consciousness separately, once relying on the paradigms, and again using the neuroscientific techniques (e.g., EEG, fMRI, intracranial recordings, MEG, TMS etc.) as factors (see Supplementary Figure S5, for the distribution of experiments according to neuroscientific techniques). RPT was excluded from the analysis of state consciousness and IIT was excluded from the analysis of content consciousness, due to not having sufficient number of papers. For both post-hoc analyses within studies focusing on state-consciousness, the accuracy of the classifier was not significantly above chance (see Supplementary Figure S6e-h). For content consciousness studies, classification was above chance for the classifier that predicted theory support based on neuroscientific techniques (accuracy = 62.71%, *t*(290) = 2.96, *p*= 0.007, with a chance level at 56.7%; see Supplementary Figure S6a-b) and did not survive correction for multiple comparisons for the classifier trained on experimental paradigms (accuracy = 61 %, t(290) = 2.01, p = .061; see Supplementary Figure S6c-d). None of the factors in these analyses was identified as important, suggesting that it is the combination of factors, rather than any specific factor, that drove the classification (see Supplementary Tables 7 and 8; also confirmed by the aggregated results over all classifiers used for the sensitivity analysis of these models). This suggests that generally speaking, when inspecting content and state studies separately, the data does not indicate a strong methodological bias towards one of the theories, either because it does not exist, or due to insufficient amount of data to detect a bias.

### Neural correlates of consciousness and their interpretation in light of the theories

Aggregating neural findings reported in the experiments in our database reveals a remarkable heterogeneity of findings, which by itself is not compatible with the predictions of any of the theories (that is, none of the theories would predict such a vast neural activation as a marker of consciousness). At the anatomical level, a map of all reported findings seems to suggest that almost the entire brain has been implicated in conscious perception (Figure 6a). Yet when the experiments are divided according to the theory they support, four completely different patterns emerge, that by and large are well aligned with the predictions of each of the theories. This raises the concern for a bias, where each theory confirms itself, while ignoring other findings that are incompatible with its predictions.

**Figure 6.**
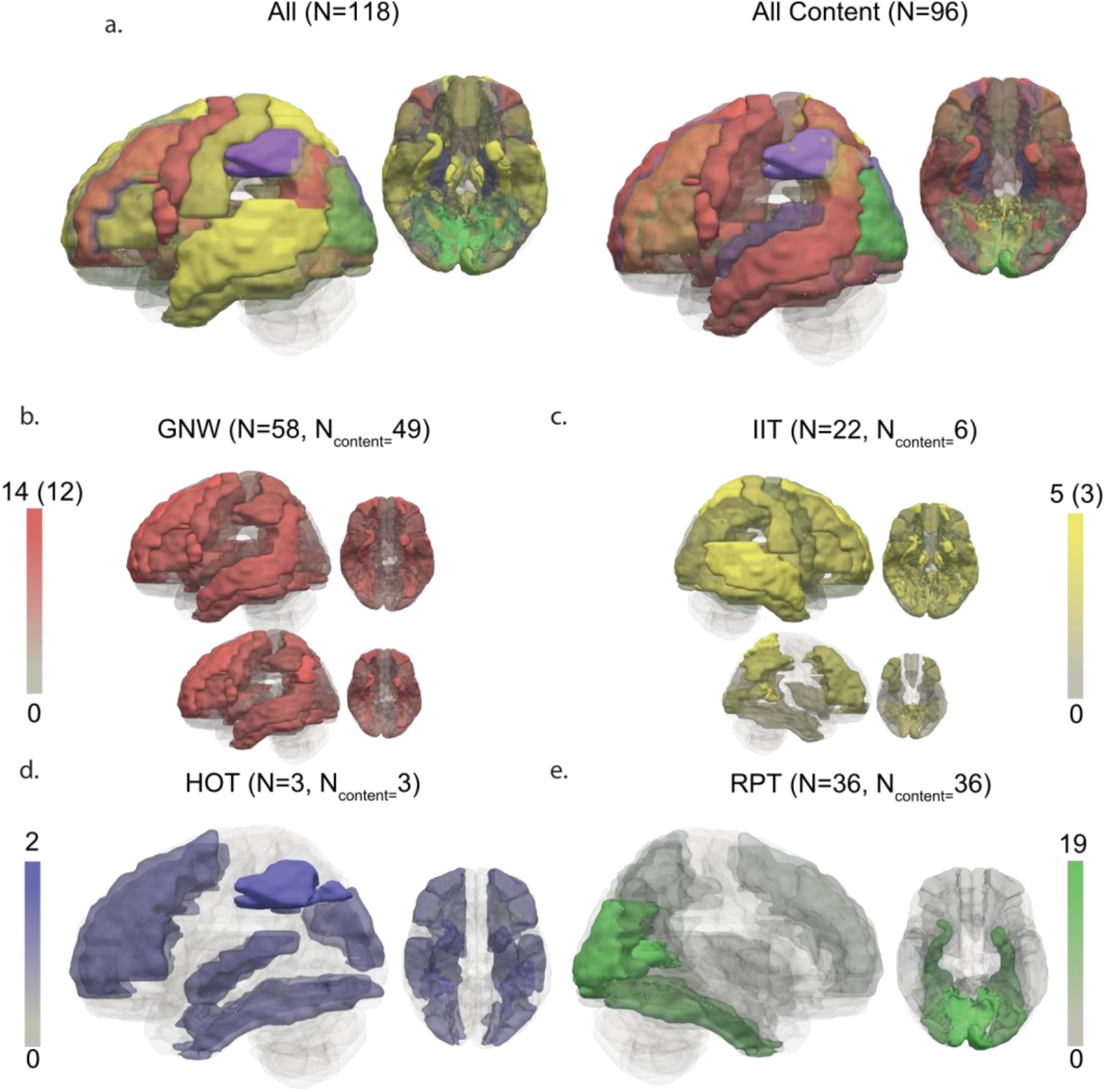
Spatial findings. Panel a: an overlay of fMRI findings reported in all experiments in the database (left) and all experiments focusing on content consciousness (right), using the AAL3 atlas^51^. Red, yellow, blue, and green activations represent experiments supporting GNW, IIT, HOT, and RPT, respectively. The intensity of the color of each activation indicates the relative frequency of experiments reporting activations in each brain area (see Supplementary Figures S7 for a similar figure, in which findings of all experiments in the database are overlaid irrespective of theory, and Supplementary Figure 8 for a figure comparing activations found in report and no-report paradigms, revealing interesting differences between the two). Panels b-e: the same findings, separately presented for experiments supporting GNW, IIT, HOT and RPT respectively, using the same color coding. Panels b-c also depict findings reported only in experiments focusing on content consciousness, presented below the maps for all experiments (in panels d-e, for HOT and RPT, all experiments focused on content consciousness to begin with). The color scales specify the number of experiments reporting activations in the same area, ranging from 0 to the maximal number of such experiments for each theory.

In the temporal domain, a similar picture emerges; when all the data from experiments using EEG, iEEG or MEG are examined together, irrespective of the supported theory, the only clear pattern is the great variability of the data, with no clear-cut answer about the timing of the NCC (Figure 7a). But when divided to theories, the reported timings (e.g., latency, peak, or pre-defined time-window; for all the above we focused on the earliest time point reported) reveal later NCCs in experiments supporting GNW (M=290.26ms, SD=142.64; Figure 7b) compared with experiments supporting RPT (M=245.65 ms, SD=110.6; Figure 7c. t(178) = 2.33, p = .034, comparing reported components). Interestingly, limiting the observations to theory-driven experiments only (i.e., excluding papers that interpret their findings post-hoc, or only mention the theories; Figure 7d-e) reveals a more dichotomous picture of NCC timing, strongly aligned with the predictions of the specific theory targeted by each study (GNW: M=340.17ms, SD=170.58; RPT: M=227.85, SD=75.38; t(55)= 3.1, p = .007). It appears as if experiments that only interpret their findings post-hoc in light of the theories find both early and late components, while researchers who set-out to test a prediction of a theory find evidence that fits well with the tested predictions. Another notable observation pertains to the variability of reported timings within a given EEG/MEG component, being most pronounced in post-hoc interpreted experiments. That is, the reported timing of the very same component (e.g., VAN ^52^ or P3 ^53^) substantially varies between studies, in the magnitude of hundreds of milliseconds (VAN range: 130-460 ms, M = 251.48 ms, SD = 67.88; P3 range: 130-908 ms, M = 446.39ms, SD = 108.1), which is not in line with the explicit predictions of the theories themselves^3,54^. This raises the concern that researchers have a large degree of freedom when interpreting and reporting their findings.

**Figure 7.**
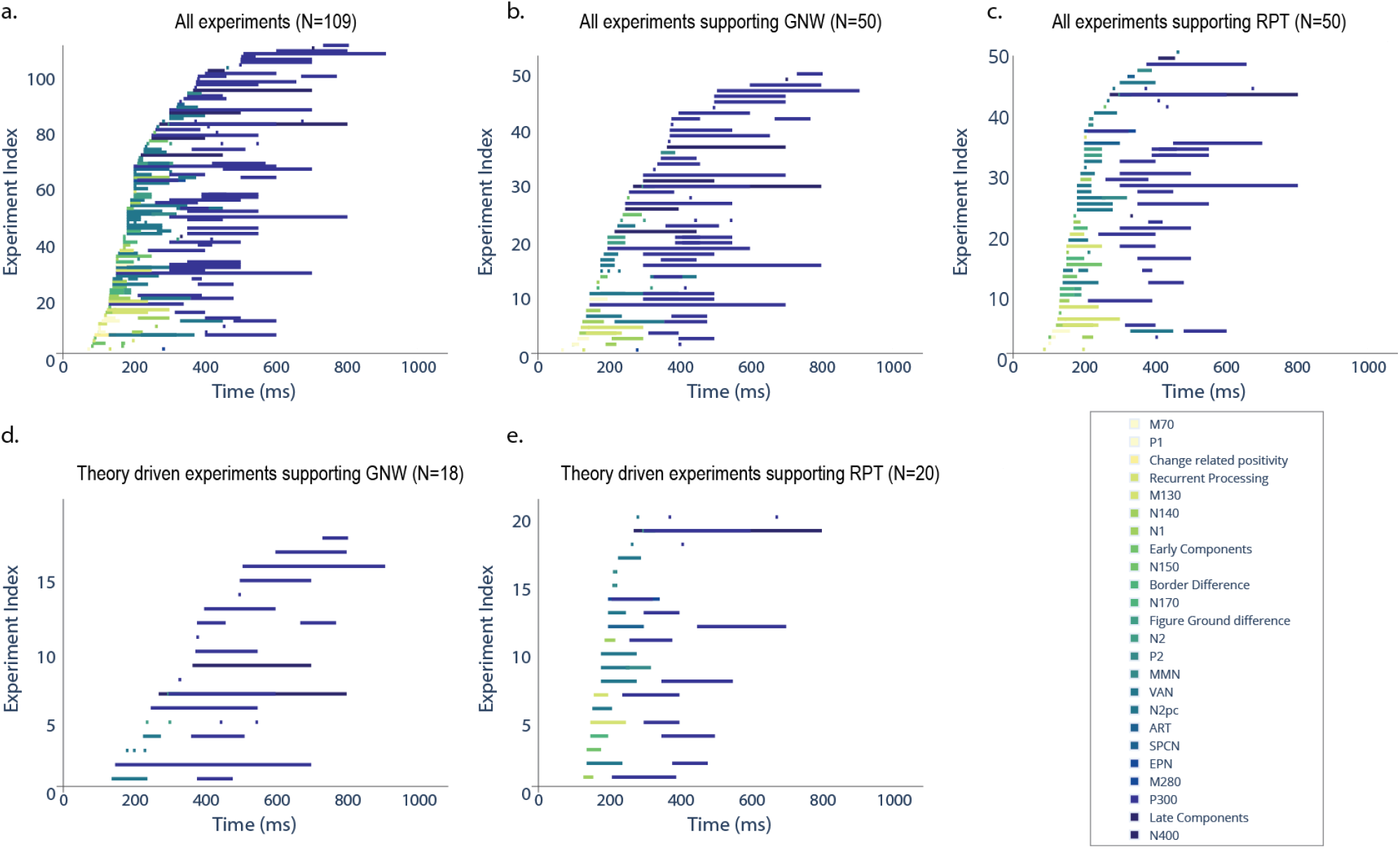
Temporal findings of EEG, iEEG and MEG components that were reported in the experiments in the database. Findings are sorted based on the earliest reported timepoint for each component, and plotted together, with the y-axis referring to an arbitrary index number of the different experiments generated for each plot. Each horizontal line represents a specific component, colored according to its classification by the authors (see the legend), with darker colors indicating later components. Components for which only a single time point was reported (e.g., peak), are represented by dots. a: temporal findings across all experiments. b: temporal findings in experiments supporting GNW only. c: temporal findings in experiments supporting RPT. Panel d, e: Similar to Panels b and c, except that the experiments were filtered to include only theory-driven studies.

## Discussion

Several key conclusions can be drawn from our analyses of these 412 experiments: First, the field seems highly skewed towards confirmatory, as opposed to disconfirmatory, evidence^55^, which might explain the failure to exclude theories and converge on an accepted, or at least widely favored, account. This effect is relatively stable over time. Second, theory-driven studies, aimed at testing the predictions of the theories, are rather scarce, and even rarer are studies testing more than one theory, or pitting theories against each other – only 7% of the experiments directly compared two or more theories’ predictions. Though there seems to be an increasing number of experiments that test predictions a-priori in recent years, a large number of studies continue to interpret their findings post-hoc in light of the theories. Third, a close relation was found between methodological choices made by researchers and the theoretical interpretations of their findings^56^. That is, based only on some methodological choices of the researchers (e.g., using report vs. no-report paradigms, or studying content vs. state consciousness), we could predict if the experiment will end up supporting each of the theories. This might represent the different emphasis put by the different theories on different aspects of consciousness. These three key findings, taken together, help explain why theories of consciousness continue to co-evolve with little impact on each other, and highlight the need for more cross-talk between the theories and more inclusive choices of paradigms testing each one of them. Notably, all of our results can be regenerated in an open-access website (http://ContrastDB.tau.ac.il/) in which all parameters extracted from the studies in our database are available, together with online analytic tools. This unique database further allows researchers to query and analyze the data in novel ways, which might unravel additional trends in the study of consciousness. It also constitutes a powerful tool for finding relevant studies based on parameters of interest (e.g., all fMRI studies supporting GNW, or all studies with a patient population using EEG).

Our analysis of the database highlights the way theories of consciousness have evolved so far: even when one theory gains accumulating support, opposing theories remain unaffected. On the contrary, all four theories seem to progress in parallel and continue to grow throughout the years. This observation is further strengthened by the relative scarcity of experiments that challenge a theory (15%). This potential confirmation bias is not unique to the field of consciousness; it has been documented decades ago^57^, and widely discussed since in various areas of psychological research^55,58^. This might be regarded as a part of the natural progression of an emerging field of research; it typically starts from a more bottom-up approach, mostly focusing on accumulation of evidence, and – as theories start to emerge – the data is interpreted in light of the theories. Only later on the competing theories are critically tested in a top-down manner. Based on our analysis, the field of consciousness studies seems to still be well within the initial state, as the predominance of supporting experiments over challenging ones is stable over time. Here, we suggest that with the maturation of the field – both theoretically^3,4^ and methodologically^59^ – the time has come for it to transition into a more critical phase of directly testing and possibly eliminating theories, as a necessary means for making progress^60^. This could be achieved by identifying opposing predictions of two or more theories and using a design that experimentally pits them against each other^61^.

This shift seems even more important when taking into account the gap between how the neural findings look when aggregated across all studies, as opposed to their patterns when divided based on the theories they support (Figures 6 & 7). Despite the heterogeneous pattern observed when all experiments are collapsed together, each theory seems to find evidence for its specific predictions. This might reflect a different focus on certain aspects of consciousness by each theory. Conversely, this might stem from emphasizing specific results that fit the predictions while overlooking others, or from refraining from testing alternative interpretations to begin with. In addition, our analyses highlight a substantial amount of flexibility in interpretation, e.g., the timing of NCC measured by EEG and MEG. This is yet another form of a confirmation bias, where findings are interpreted in light of the expectations of the experimenters^62^.

Another clear trend we found relates to the strong focus on content consciousness. This accords with the general strategy of searching for the neural correlates of a specific phenomenal aspect of an experience as an effective means to neuroscientifically study consciousness ^63^. The strategy typically relies on contrasting between brain activity when subjects are conscious of certain content, and when they are unaware of the same content (or are aware of some other content) (e.g. ^64^, but see ^59,65^). In contrast, studies focusing on state consciousness target differences between conditions in which subjects are in an overall state of unconsciousness (e.g., dreamless sleep, anesthetized, suffering from a disorder of consciousness, etc.) as opposed to a conscious one (e.g., wakefulness, dreaming, recovery from a disorder of consciousness, etc.). Notably, both of these approaches have been criticized based on different grounds: studying *content* consciousness usually involves keeping the state of consciousness constant, thereby neglecting the neural processes necessary for being in that state to begin with ^66^. Also, such studies typically assume that the contrast between perceived and not-perceived stimuli indeed distills the correlates of conscious perception, while often failing to account for confounding processes that either pertain to the prerequisites or the consequences of consciousness ^59,67^. On the other hand, studying *state* consciousness inherently implies a difference between experiencing some contents vs. experiencing no contents at all^66^. In addition, the contrast between different states of consciousness typically involves several uncontrolled, or difficult to control, confounding variables (e.g., overall arousal, attention, etc.). These complexities, in turn, limit the ability to detect mechanisms that are uniquely correlated with consciousness per se ^63^. Thus, a refinement of predictions to address both content and state, along with development of novel ways to test existing predictions while avoiding confounding factors, will be necessary for achieving progress. These directions are important because all four theories make claims about both content and state consciousness ^4,32,63^, and should indeed account for both aspects to provide a comprehensive explanation of consciousness^4,32,66^.

Our review was epistemic and descriptive in nature, taking a hands-off neutral approach, accepting the interpretations made by authors at face value. Future efforts could complement this database with a meta-analyses aimed at testing the reliability and validity of findings, and exploring in more detail potential differences between the results themselves and the authors’ interpretations of such results. Such meta-analyses could also include additional papers that were not the focus of the current study (e.g., NCC papers that refer to other theories, or not referring to any theory), explore other search strategies, and statistically examine the meta-analytic reliability of findings across studies. This was not our goal here (though, our open-access database and website could be used as a first step for doing so). The data extraction procedure in our case is solely based on the way the authors themselves interpreted their results, so that even if we found the interpretation inaccurate/erroneous (e.g., claiming that GNW is supported by very early activations, or that RPT is supported by frontal findings), the paper was still included in the analysis as is. We also took a neutral approach to how the theories have evolved over the years (e.g., IIT once suggested that frontal areas could interact with posterior ones to generate conscious perception^68^ but claims differently today^28,40^; and GNW once treated P300 as a clear marker for consciousness^69,70^ and is more hesitant in doing so today^25,32^). Thus, findings that were once taken as evidence in favor of a theory, might actually be taken today as evidence against it, and this would not be reflected in our analysis. Our review also calls for a better differentiation between core predictions made by a theory, and auxiliary ones that are less diagnostic for testing it (e.g., the change with respect to P300 described above is relevant for an auxiliary prediction by GNW, not to a core hypothesis). Since such a dissociation is not clear in existing literature – and in the papers we examined – we were unable to make that distinction in our database. We accordingly invite theory leaders, as well as the field as a whole, to explicitly pinpoint core and auxiliary hypotheses. This could then be integrated into our open database, propelling further insights into the theories and the progression of the field as a whole.

To conclude, although the field of consciousness studies has been enjoying great proliferation in the last decades, the impressive body of literature accumulated has yet to converge into a widely-accepted theory. As the field matures, such a convergence is more likely, yet to reach that state, it seems that several key steps should be taken. First, future studies should focus more on testing theories of consciousness, rather than looking for confirmatory evidence, or relying on post-hoc interpretations that allows them to co-evolve without convergence and/or elimination of some theories. The confirmation biases we found across theories, as well as the relatively low number of studies that set-out a-priori to test the theories, highlight the need to do so. Second, it will be important to expand the type of experiments testing each theory^66^, for example examining both state and content consciousness, over different populations, measures, and tasks. Given that all theories make claims relevant to both content and state consciousness^32,35,71,72^, it would be advisable to avoid limiting investigations to one type. Similarly, all of the leading theories considered here attempt to explain the neural basis of consciousness broadly speaking and should therefore be testable using a wide-variety of neuroscientific measures, manipulations of consciousness, and report and no-report tasks. Despite some of the trends and potential biases uncovered by the analyses reported here, we remain optimistic that honest attempts for collaborations, and testing opposing predictions in an unbiased manner will ultimately lead to theory refinement, elimination, and convergence in the quest to understand consciousness from a neuroscientific perspective.

## Methods

### Creation of paper database

The database was created following predefined criteria, that are reported below in line with the PRISMA 2009 guidelines ^41^ concerning inclusion and exclusion criteria, search strategy, and data collection procedure.

### Inclusions and exclusion criteria

The database includes studies that conform with the following criteria: (C1) the study reports empirical results published in a peer-reviewed journal, written in English; (C2) the study pertains to the neural correlates of consciousness; (C3) at least some of the findings reported in the study were interpreted in light of one or more of an NCC prediction of the four theories of consciousness reviewed here (GNW, HOT, IIT, RPT); and (C4) the study was conducted using a neuroscientific technique. Papers that were first detected in the initial search sweep (see below) but did not meet the above criteria, were excluded. Screening of these papers in light of these criteria was done by I.Y., and papers for which there was a dilemma concerning one or more of the criteria, were further screened by L.M^a^.

### Search strategy

Scopus electronic database (up to 22/10/2019) was searched using two separate search strategies: (1) **Topic search**, using the name of each of the four theories as part of the topic / abstract / keywords of papers in the database; (2) **Citation search**, where we first identified three key papers for each theory (for a list of the key papers, see Supplementary Table S1). The key papers were chosen in a joint discussion between I.Y., L.M^a^, L.M^b^, and M.P. Then, all papers citing one or more of these twelve papers (three papers times four theories) were selected.

No constraints were enforced on the initial Topic and Citation searches. Then, the following filters were applied: (i) exclude non-English records; (ii) include only records with ‘document type’ = ‘Document’; (iii) include only records classified in the Scopus database as belonging to the categories: ‘Neuroscience’ / ‘Psychology’ / ‘Multidisciplinary’ (for a detailed description of the distribution of papers in both included and excluded categories, see Supplementary Table S2). For each of the theories, the search was conducted using the query: TITLE-ABS-KEY (“X”), where “X” denotes the theory specific topic keywords. The same query was used for the citation searches to find the entry of each key paper, from which we collected the papers that cited them using the Scopus interface. Then, the above-mentioned filters were applied: (i) LIMIT-TO (LANGUAGE, “English”) ; (ii) LIMIT-TO (DOCTYPE, “ar”) ; (iii) LIMIT-TO (SUBJAREA, “NEUR”) OR LIMIT-TO (SUBJAREA, “PSYC”) OR LIMIT-TO (SUBJAREA, “MULT”). Lastly, papers that include one or more neuroscientific technique keywords (EEG, Imaging, fMRI, ERP, Neuroimaging, TMS, MEG, intracranial, PET, ECoG, Electrophysiology, single units, iEEG and multi-units) in their abstract were detected using an in-house script we developed.

Figure 1a (see Results above) presents the process of article selection and screening, and Figure 1b-c describes the division of papers to search strategies, before the screening and following it. Out of 6938 records identified in the initial database searching, 2845 unique papers passed the first screening stage (constraints i – iii detailed above, screened serially on the database). 187 papers were excluded as they were not written in English (i); 2292 records were filtered out since they were not of the “Document” type (ii); and 947 did not belong to one of the predefined categories (iii). Enforcing the last constrain of using a neuroscientific technique resulted in excluding 1961 papers. The remaining 884^c^ unique papers were assessed for eligibility by a close inspection of their full-text articles conducted by I.Y., with the supervision of L.M^a^, and further discussion of unresolved issues with the collaborators L.M^b^ and M.P, until a consensus was reached. In this process of close inspection, 519 papers were excluded due to the following reasons: (1) not relating to consciousness studies (n = 232). These papers focused on other phenomena, like working memory or intelligence or attention, and mentioned one or more of the theories concerning those phenomena, and not consciousness (see Supplementary Figure S1 and Supplementary Table S2 for the distribution of excluded papers into main fields of research); (2) mentioning one or more of the theories, but did not interpret their findings with respect to any of them (n = 190); (3) Reviews (n = 89); (4) Meta-analyses (n = 5) ; (5) Behavioral studies that still mentioned a neuroscientific technique in their abstract, despite not using it (n = 3). This process yielded the final database, which included 365 papers, reporting 412 experiments, all of which were classified according to our predefined parameters of interest.

### Data Collection

A custom data extraction sheet was developed. For each paper and nested experiments, we automatically extracted the following metadata using Scopus’ web interface: title, DOI, authors, affiliations, author keywords, index keywords, source title, publisher, funding, references, abstract, and number of citations. Then, information about our predefined parameters of interest was extracted manually: main experimental paradigm, specific experimental paradigm, indicator whether the experiment used report / no-report paradigm, indicator whether the experiment studies content / state consciousness, sample type, total sample size, sample size of included subjects, task description, task type, stimuli categories, stimuli description, stimuli modality, stimuli duration, stimuli contrast, consciousness measure taken, consciousness measure type, consciousness measure description, neuroscientific technique, a summary of the findings, findings coded as tags, dependent measures taken, interpretation regarding each theory, two Boolean parameters indicating whether the experiment is an internal replication and whether it was theory-driven, and lastly the ALL3 label for effects reported in fMRI studies (for a full description of the extraction sheet see Supplementary Table S4). The extraction of these parameters was done by I.Y. Any dilemma concerning the classification of parameters or inference regarding the interpretations of the authors were resolved by L.M^a^.

Critically, the parameters of interest were extracted based on how they were presented in the original paper. That is, no attempt was made to re-interpret or test the statistical reliability of the original findings and consequent interpretations or to critically assess the strength of the experimental design. Choosing to take the reported findings and interpretations at face value was in line with the goals of this quantitative review; as opposed to a meta-analysis, where one wishes to assess the strength and reliability of certain effects ^73^, our goal was to characterize the field of consciousness studies: *what claims are being made and based on what types of findings and paradigms*. Moreover, sticking to the original interpretations of the authors allowed us to keep this review objective and uncontaminated by our subjective points of view.

### Neural Data Extraction

An AAL3 ^51^ label was encoded for each candidate NCC brain area found in an fMRI experiment, according to the reported coordinates. A script^74^ (label4MNI) written in R was used to map MNI coordinates into the respective AAL labels. When Talaraich (TAL) coordinates were reported, they first were transformed to MNI coordinated using the MNI <-> TAL online converter^75^; based on the mapping reported in ^76^), and then AAL labels were extracted using the label4MNI script. In cases where no coordinates were reported whatsoever, which was the case in most experiments using functional localizers, Neurosynth ^77^ was used to extract the MNI coordinates showing the greatest fit for the reported brain area according to the label used by the original authors. Then, for each of the theories, nii masks for specific brain areas were generated using Matlab, by uniting the masks of all AAL3 labels extracted from papers supporting the theory. The masks were processed to form a 3D model of the brain areas associated with each theory, using both ITK-SNAP ^78^ and Paraview ^79^, and following the procedure suggested by Madan^80^. The resulting 3D models were then overlaid on a 95% opaque AAL brain and colored according to the respective theory. The opacity and intensity of the color of each area were set according to the frequency of activations, normalized by the frequency of the most frequently reported area. Note that laterality was not encoded for the AAL3 labels.

### Open access and online website

All analysis and pre-processing codes used in this paper are shared in OSF^81^ (https://osf.io/avz8b/). Also, we made our database publicly accessible in a website we developed using the Dash Plotly^82^ framework. In the website we enable free querying of the database, and provide interactive visualization tools. By exposing the database to the community via an easy-to-use graphical interface we encourage researchers in the field to examine their meta-questions about the field, possibly leading to new lines of informed research about the theories of consciousness. Moreover, in addition to the dynamic querying interface which facilitates cutting the data according to flexible conditions, the website includes interactive graphs describing sociological information about the field and enables retrieving papers of interest according different cuts of the data.

### Data analysis

Data analysis was performed using R ^83^ and Python ^84^. It included descriptive reports of the distributions of the extracted parameters, as well as statistical analysis of the uniformity of these distributions. Importantly, the latter was performed including only theories supported by at least 20 experiments. Similarly, only parameters with 10 or more observations (e.g., methods reported in at least 10 experiments) were included.

The uniformity of the distributions was either tested with a Chi-square test, or using random forest classifiers ^85^, trying to predict the outcome of an experiment based on the parameters. This was done under the assumption that above-chance classification reflects a non-uniform distribution of parameters (i.e., a specific combination of parameters is more likely to predict the support of specific theories). Random forest is a machine learning technique supporting multi-class classification by learning complex patterns in multidimensional data using multiple decision trees ^86^ and bootstrapping procedures. Here, random forest classifiers with a zero random state (acts as a seed for the random processes used to build random forest classifiers), leaving the remaining parameters at the default values of the scikit-learn python package^87^, were used to test whether specific methodological parameters reported in experiments studying consciousness can be used to predict the theoretical interpretation of its findings. The model included all extracted parameters, yet we excluded parameters due to multicollinearity (for a full list of extracted parameters, see all items on Supplementary Table S6 that are marked with an asterisk) ^43^. Specifically, we removed parameters with a variation inflation factor (VIF) of more than 5^88,89^ from the full model. For completeness, we report the results of a full model, from which no parameter was excluded, in the supplementary material (Supplementary Figure S2, and Supplementary Table S6).

The predictive power of the classifiers was assessed using a leave-one-out strategy: support/no-support for each one of the theories was iteratively predicted by the parameters of each specific experiment, yielding a 0/1 metric that was based on the success of the classifier to predict the outcome for that experiment. To allow this binary outcome, experiments that found evidence against a theory or remained neutral regarding the theory were both considered as not supporting it, without differentiating between these two cases. Then, three analyses were conducted to quantitively assess the performance of the classifier: first, a t-test was used to compare the accuracy of the classifier with the accuracy of a ‘chance-level’ model which predicts the support of each theory without taking into account any of the parameters, solely based on the frequency of the outcome in our database (i.e., how often is a specific theory supported). Thus, the chance-level model should maximize prediction accuracy given the marginal distributions of the support of each theory. In this analysis, the dependent variable was the accuracy of the two models (chance level model vs. our model, using the parameters), defined as the averaged accuracy across all 412 iterations, where in each iteration accuracy was defined as the percentile of correct classifications (out of three, as support for each one of three theories was predicted). Second, to test the robustness of the results, and control for the randomness of the random forest classifiers fitting procedure, we conducted a sensitivity analysis. Specifically, we compared the results of our main analysis detailed above with classifiers trained and tested on the same data, yet with 1000 different random states. We then tested in how many of these iterations the same results are obtained. Third, an SDT^44^ was used to further assess classification performance for each theory. Specifically, a Receiver Operating Characteristic curve (ROC) was calculated for each theory, and the respective Area Under the Curve (AUC) was reported. AUC weights both true and false-positive predictions of a classifier and is unaffected by imbalances between positive and negative cases (which makes it especially suitable for this analysis, since the support for the different theories is unequally distributed). AUC of 0.5 indicates the classification performance of a random classifier.

When the classifier showed significant performance, the importance of each feature in the model was assessed using a permutation-based importance assessment on the entire training data ^90^. This method provides a measure of how classification performance changes when the values of each parameter change (i.e., are permuted) and thus should indicate which specific parameters contributed to the observed classification performance. The Importance score for each parameter was calculated based on 5000 permutations of the values of each parameter in 1000 iterations where the model was trained with an additional, newly sampled, random parameter. Then, they were compared with the importance of the random parameter added to the training data, to evaluate the reliability of the importance assessment result. Only features with an average importance score higher than the 95% quantile of the importance of a random variable were considered important for classification. Akin to the sensitivity analysis of classifier accuracy, we ran a similar procedure for the importance scores. Here, we calculated importance scores for each of the 1000 classifiers, as described above, and aggregated the results of all classifiers together. Importance scores were calculated based on 5000 permutations of each parameter in 100 iterations with newly sampled random parameter.

All p-values reported throughout the manuscript were adjusted using false discovery rate (FDR), ^91^ to account for possible alpha inflation due to multiple comparisons.

## Acknowledgements

This project was made possible through the support of grants from Templeton World Charity Foundation Inc. (TWCF0378, TWCF0599), and the National Science Foundation (BCS-1829470). The opinions expressed in this publication are those of the authors and do not necessarily reflect the views of Templeton World Charity Foundation, Inc.

## Footnotes

a Liad Mudrik

b Lucia Melloni

c The papers extraction procedure resulted in finding 885 unique papers, yet after manual inspection one duplicate was found.

