## Supplementary Material for "The Consciousness Theories Studies (ConTraSt) database: analyzing and comparing empirical studies of consciousness theories"

### Supplementary figures

#### Supplementary Figure S1: Distribution of papers not studying consciousness

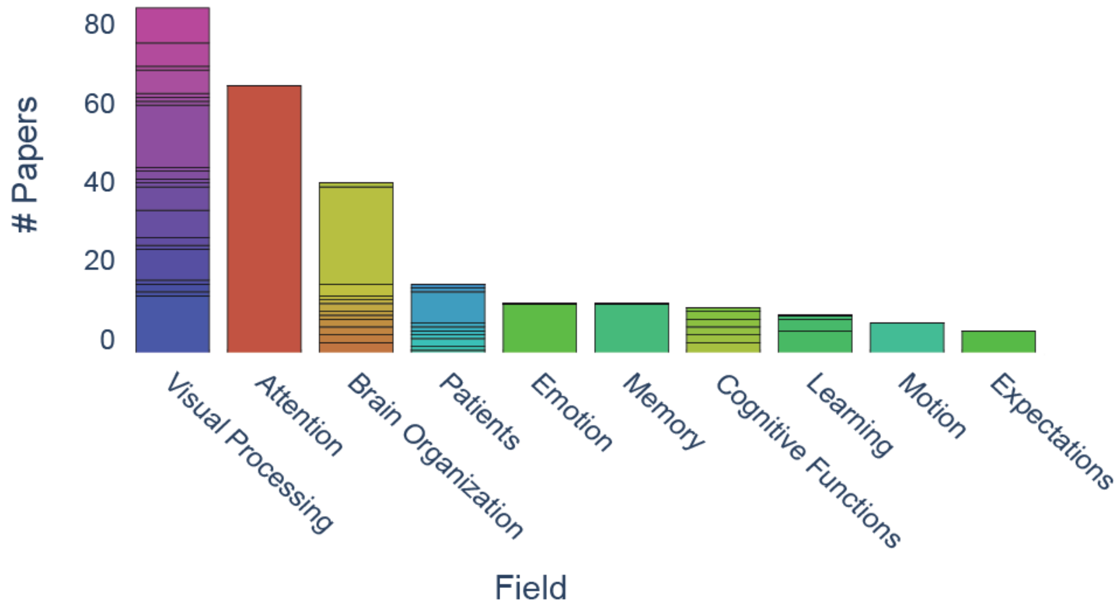

Figure S1. Distribution of papers that were excluded after close inspection due to the criterion 'not studying consciousness', according to their topics of research. See Table S3 for a detailed description of the distribution. Colors within each bar denote different topics within the general category listed on the x-axis. Fields with less than five papers were not included in the figure.

### Supplementary Figure S2: Classifier performance, full model

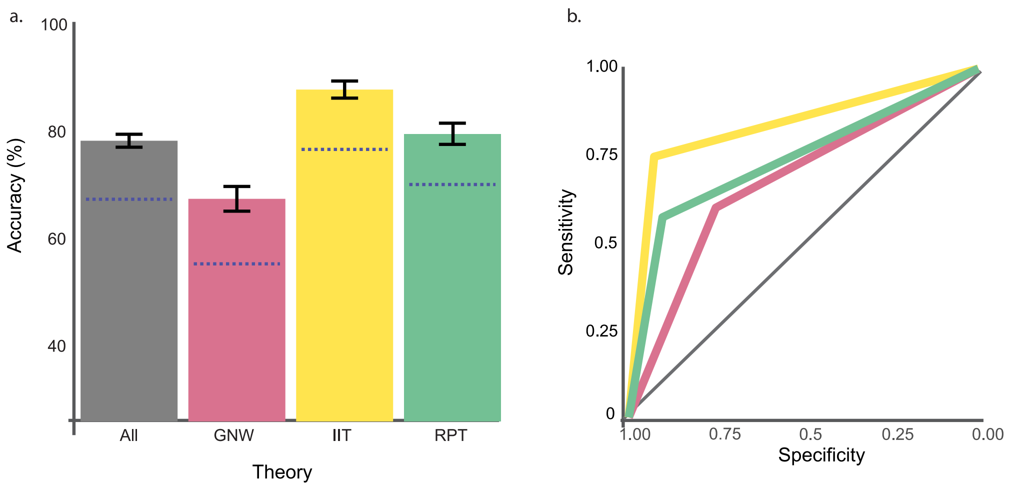

Figure S2. Analysis of the classifier's performance in predicting support for each theory, when including all parameters in the database (without exclusions due to multicollinearity). Panel a: The classifier reached accuracy of 78.56% compared with chance-level performance of 67.6% (sensitivity analysis: M = 79.47%, SD = .46%, range: 77.83%-81.67%). Panel b: classification accuracy for all three theories (gray), and each individual theory: IIT (yellow), RPT (green), and GNW (red). Panel b: ROC curves for classification performance. Each line performance for each of the theories separately; the gray diagonal line describes the performance of a random classifier. Blue dashed lines indicate chance level performance for each of the conditions.

### Supplementary Figure S3: Feature Importance, main model

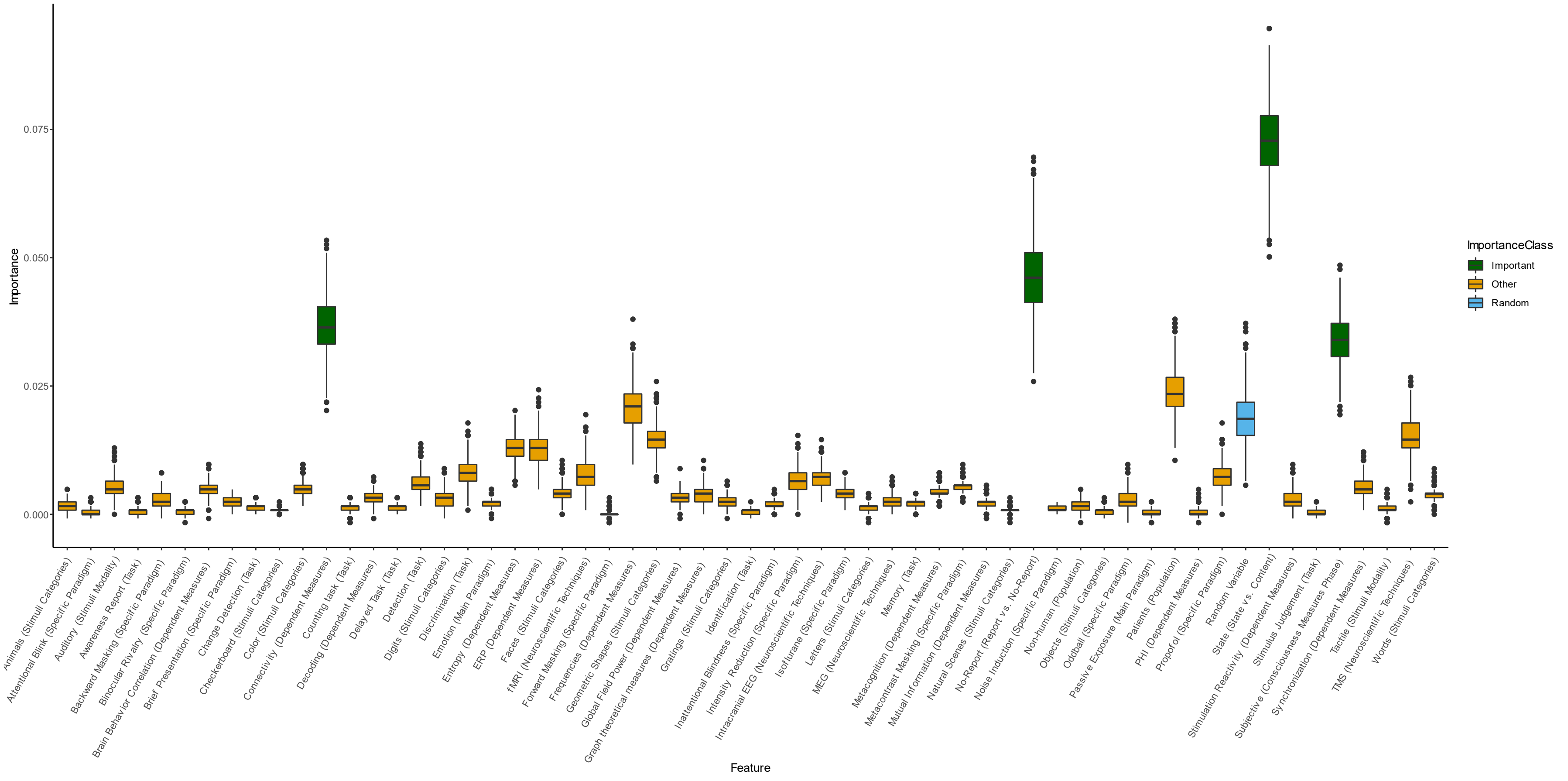

Figure S3. Feature importance analysis of the features included in the main analysis. Each box depicts the distribution of 1000 importance scores calculated using a permutation-based method with 5000 samples. Green and Orange boxes describe features with mean importance score higher and lower (respectively) than the 95$^{th}$ percentile of importance scores calculated for a random feature (Blue).

### Supplementary Figure S4: Division to paradigms

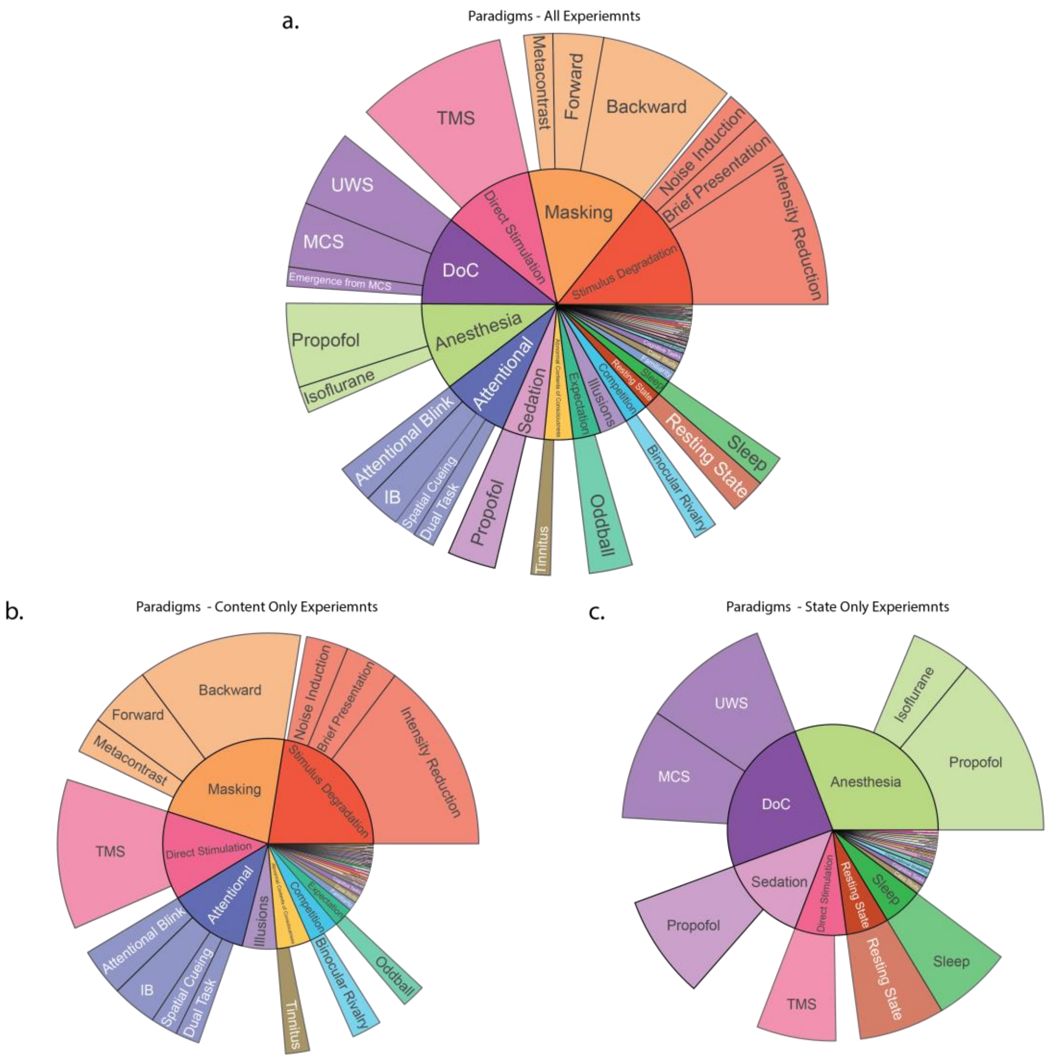

Figure S4. Panel a: distribution of experimental paradigms. The inner-circle describes the distribution of the main paradigms, and the outer circle describes the distribution of specific paradigms within each main paradigm. Panel b, c: the same as Panel a, except that the distribution is limited to experiments studying content and state consciousness, respectively. Slices with less than eight experiments were not included in the figure. Note that since experiments can use multiple paradigms, N can exceed the number of experiments in the database (412). Abbreviations: AB (Attentional Blink), BR (Binocular Rivalry), DoC (Disorders of Consciousness), IB (Inattentional Blindness), MCS (Minimal Consciousness State), TMS (Transcranial Magnetic Stimulation), UWS (Unresponsive Wakefulness State).

### Supplementary Figure S5: Division to neuroscientific techniques

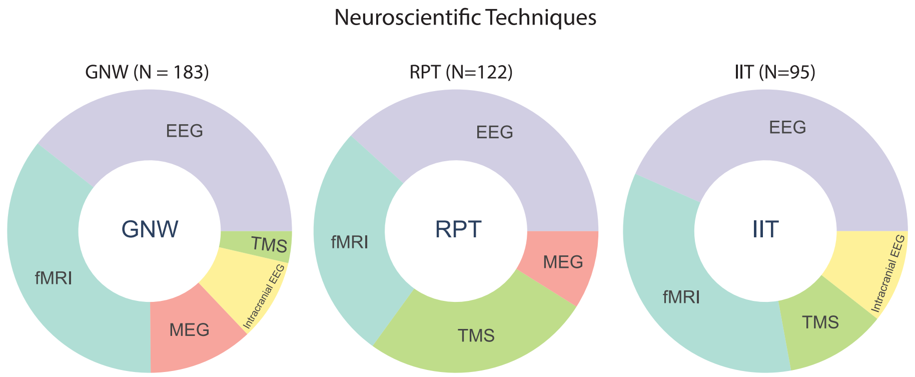

Figure S5: Distribution of experiments supporting a certain theory according to the neuroscientific technique used in each experiment. Slices with less than five experiments were not included in the figure. Abbreviations: EEG (Electroencephalogram), fMRI (Functional Magnetic Resonance imaging), MEG (Magnetoencephalography), TMS (Transcranial Magnetic Stimulation).

### Supplementary Figure S6: Classifier performance, post-hoc models

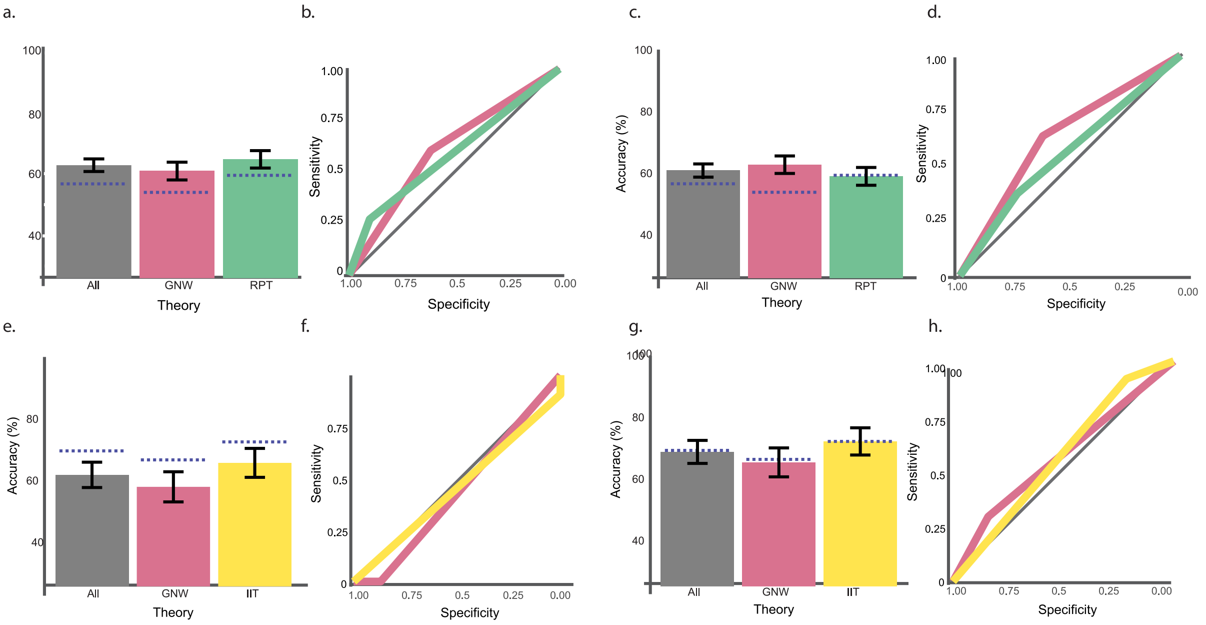

Figure S6. Analysis of the classifier's performance in predicting support for each theory, for content and state consciousness separately. Upper panel: Content consciousness: Statistical analysis showed significant accuracy for Content consciousness: accuracy of 62.71% and 61% vs. chance of 56.7% for the neuroscientific techniques (e.g., EEG, fMRI, intracranial recordings, MEG, TMS etc.) (panels a-b), and paradigms (panels c-d) models, yet the latter effect did not survive correction for multiple comparisons, t(290) = 2.01, p = 0.45 uncorrected, p = 0.063 corrected, and did not show consistent above chance performance according to sensitivity analysis (M = 61.05%, SD = 1.09%, range: 55.33%-63.57%) in contrast to the neuroscientific techniques model, t(290) = 2.01, p = .007 corrected, and more consistent sensitivity analysis results (M = 62.87%, SD = 0.2%, range: 61.68%-63.06%); Lower panel: State consciousness: accuracy was below chance, 62.13% and vs. 69.9% (sensitivity analyses results: M = 65.85%, SD = 2.19%, range: 61.65%-69.9% and M = 69.43%, SD = .99%, range: 66.02%-72.33%, respectively), for neuroscientific techniques (panels e-f) and paradigms (panels g-h) models. All p-values for the state consciousness classifiers were higher than 0.05 before correction). For each classifier, both classification accuracy (panels a,c,e,g) and ROC curves (panels b,d,f,h) are depicted for all three theories (gray), and each individual theory: IIT (yellow), RPT (green), and GNW (red), based on the paradigm used in the experiment. The gray diagonal line describes the performance of a random classifier. Blue dashed lines indicate chance level performance for each of the conditions.

**Supplementary Figure S7: Spatial findings across theories**

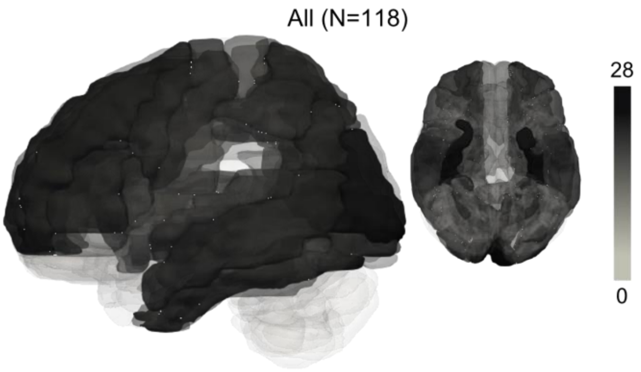

Figure S7: An overlay of fMRI findings reported in all experiments in the database, using the AAL3 atlas[1]. Darker colors indicate the frequency of frequency of the activation in each brain area collapsed across the theories. The color scale indicates the association between the color of each brain area and the number of experiments reporting activations in this area

.

**Supplementary Figure S8: Spatial findings: Report vs. No-Report within content consciousness studies**

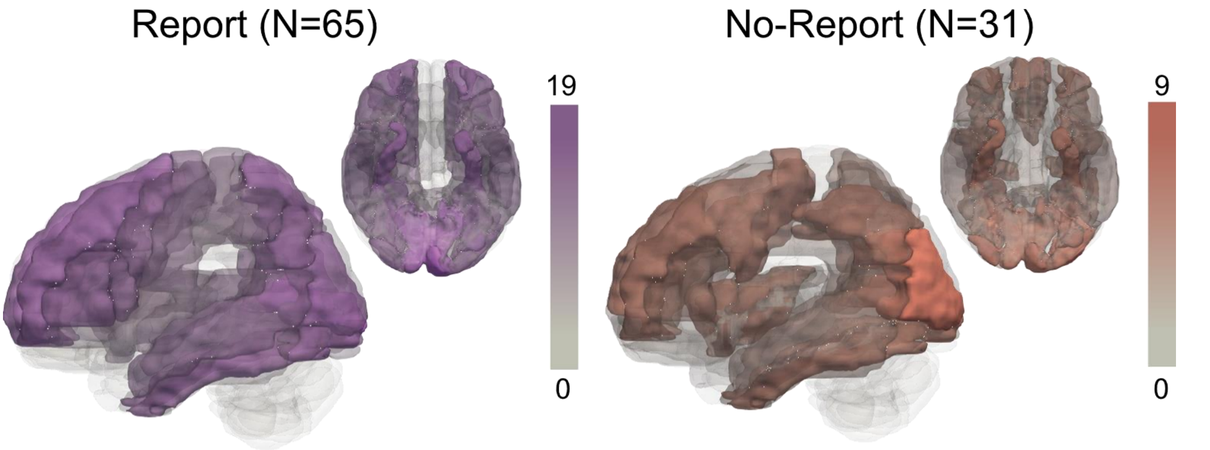

Figure S8: An overlay of fMRI findings reported in experiments focusing on content consciousness, divided to experiment that used report (left panel) or no-report (right panel) paradigms. Darker colors indicate the frequency of frequency of the activation in each brain area. The color scale indicates the association between the color of each brain area and the number of experiments reporting activations in this area.

**Supplementary Figure S9: Division to Modalities**

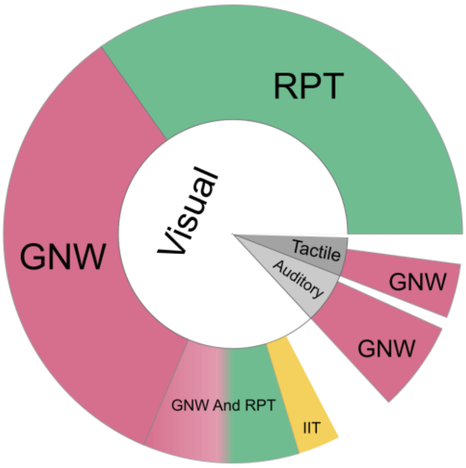

Figure S9: Stimulus modality distribution over 309 experiments focusing on content consciousness (inner circle), and division to theories within each modality (outer circle). Slices with less than five experiments were not included.

### Supplementary tables

#### Supplementary Table S1: Search strategies

| | GNW | HOT | IIT | RPT |
|---|---|---|---|---|
| Topic keywords | "global neuronal workspace" | "higher order thought" | "integrated information theory" | "recurrent processing" |
| 1st key paper | Dehaene, S., & Naccache, L, 2001, *Towards a cognitive neuroscience of consciousness basic evidence and a workspace framework*, Cognition | Rosenthal, D. M., 1993, *Higher-order thoughts and the appendage theory of consciousness*. Philosophical Psychology | Tononi, G., 2004, *An information integration theory of consciousness*, BMC neuroscience | Lamme, V. A., & Roelfsema, P. R., 2000, *The distinct modes of vision offered by feedforward and recurrent processing*, Trends in neurosciences |
| 2nd key paper | Dehaene, S., et al., 2006, *Conscious, preconscious, and subliminal processing: a testable taxonomy*, Trends in cognitive sciences | Cleermans, A, 2007, Consciousness: *the radical plasticity thesis*, Progress in brain research | Tononi, G., 2011, *The Integrated Information Theory of Consciousness: An Updated Account*, Archives italiennes de biologie | Supèr, H., Spekreijse, H., & Lamme, V. A., 2001, *Two distinct modes of sensory processing observed in monkey primary visual cortex (V1)*, Nature neuroscience |
| 3rd key paper | Dehaene, S., & Changeux, J. P., 2011, *Experimental and Theoretical Approaches to Conscious Processing*, Neuron | Lau, H., & Rosenthal, D., 2011, *Empirical support for higher-order theories of conscious awareness*, Trends in cognitive sciences | Tononi, G., et al., 2016, *Integrated information theory: from consciousness to its physical substrate*, Nature Reviews Neuroscience | Lamme, V. A., 2006, *Towards a true neural stance on consciousness*, Trends in cognitive sciences |

*Table S1. Upper row: the specific keywords used for the topic search; Lower three rows: the key papers of each theory used for the citation search.*

### Supplementary Table S2: Division of excluded papers to fields

| Field | Subfield | Amount |
|---|---|---|
| Active Sensing | Active Sensing | 1 |
| Alertness | Alertness | 1 |
| Attention | Attention | 69 |
| Auditory Processing | Auditory Processing | 2 |
| Auditory Processing | Detection | 1 |
| Auditory Processing | Expectations | 1 |
| Brain Organization | Brain Organization | 3 |
| Brain Organization | Calcium Imaging | 2 |
| Brain Organization | DTI | 2 |
| Brain Organization | Effective Connectivity | 2 |
| Brain Organization | Plasticity | 1 |
| Brain Organization | Prefrontal Cortex | 1 |
| Brain Organization | Simulations | 2 |
| Brain Organization | Structural | 2 |
| Brain Organization | Task related | 1 |
| Brain Organization | Task | 3 |
| Brain Organization | Resting state | 25 |
| Cognitive Control | Cognitive Control | 2 |
| Cognitive Functions | Cognitive Functions | 3 |
| Cognitive Functions | Mental Rotation | 2 |
| Cognitive Functions | Reasoning | 3 |
| Confidence | Confidence | 1 |
| Conscious Effort | Conscious Effort | 1 |
| Conscious effort | Conscious effort | 1 |
| Decision Making | Decision Making | 3 |
| Deviant Detection | Auditory odd-ball | 3 |
| Emotion | Emotion | 13 |
| Expectations | Expectations | 6 |
| Free Will | Free Will | 1 |
| Gestalt | Gestalt | 1 |
| IQ | IQ | 3 |
| Implicit Learning | Implicit Learning | 3 |
| Brain Synchronization | Brain Synchronization | 1 |
| Language | Language | 2 |
| Learning | Learning | 6 |

| | | |
|---|---|---|
| Learning | Incidental | 1 |
| Learning | Implicit Learning | 3 |
| Memory | Memory | 13 |
| Methodological Paper | Methodological Paper | 1 |
| Mindfulness | Mindfulness | 1 |
| Motion Processing | Motion Processing | 8 |
| Motivation | Motivation | 1 |
| Patients | Alzheimer's Disease | 1 |
| Patients | Amyotrophic Lateral Sclerosis | 1 |
| Patients | Autism Spectrum Disorder | 2 |
| Patients | Cerebral Small Vessel Disease | 1 |
| Patients | Epilepsy | 1 |
| Patients | Near-Electrocerebral Silence | 1 |
| Patients | Pervasive Developmental Disorder | 1 |
| Patients | Schizophrenia | 8 |
| Patients | Visual Form Agnosia | 1 |
| Patients | Unilateral Sturge-Weber syndrome | 1 |
| Simulations | Simulations | 1 |
| Sleep | Sleep | 2 |
| Social Psychology | Social Psychology | 1 |
| Somatosensory Processing | Somatosensory Processing | 2 |
| Top-Down Control | Top-Down Control | 4 |
| Visual Processing | Visual Processing | 15 |
| Visual Processing | Colour | 1 |
| Visual Processing | Context | 2 |
| Visual Processing | Contrast | 1 |
| Visual Processing | Faces | 8 |
| Visual Processing | Familiarity | 1 |
| Visual Processing | Feature Integration | 2 |
| Visual Processing | Figure-Ground | 7 |
| Visual Processing | Gestalt | 6 |
| Visual Processing | Gist | 1 |
| Visual Processing | Grouping | 1 |
| Visual Processing | Movie | 2 |
| Visual Processing | Natural Images | 1 |
| Visual Processing | Objects | 16 |
| Visual Processing | Occlusion | 1 |
| Visual Processing | Periodic Stimulation | 1 |
| Visual Processing | Recognition | 1 |

| | | |
|---|---|---|
| Visual Processing | Scenes | 6 |
| Visual Processing | Segmentation | 1 |
| Visual Processing | Shapes | 6 |
| Visual Processing | Words | 9 |
| Visuomotor Integration | Visuomotor Integration | 1 |
| Volition | Volition | 1 |

*Table S2. The number of papers that were included due to 'not studying consciousness' criterion split by subfields. The fields are sorted alphabetically.*

### Supplementary Table S3: Included and excluded categories

| Included | | Excluded | |
|---|---|---|---|
| **Category** | **Amount** | **Category** | **Amount** |
| Neuroscience | 2478 | Medicine | 828 |
| Psychology | 1748 | Arts and Humanities | 811 |
| Multidisciplinary | 188 | Biochemistry, Genetics and Molecular Biology | 580 |
| | | Agricultural and Biological Sciences | 466 |
| | | Computer Science | 406 |
| | | Social Sciences | 263 |
| | | Mathematics | 123 |
| | | Physics and Astronomy | 97 |
| | | Health Professions | 80 |
| | | Immunology and Microbiology | 64 |
| | | Environmental Science | 52 |
| | | Engineering | 51 |
| | | Chemistry | 19 |
| | | Business, Management and Accounting | 16 |
| | | Materials Science | 15 |
| | | Pharmacology, Toxicology and Pharmaceutics | 15 |
| | | Nursing | 12 |
| | | Decision Sciences | 9 |
| | | Economics, Econometrics and Finance | 7 |
| | | Chemical Engineering | 4 |
| | | Energy | 2 |
| | | Earth and Planetary Sciences | 1 |

*Table S3. The number of papers that were included and excluded according to the "category" criteria (iii). The original classification to categories was given by the Scopus database.*

### Supplementary Table S4: Extraction Sheet

| Parameter | Description | Valid Values | Notes | Encoding Format (per experiment) |
|---|---|---|---|---|
| *Paper* | | | | |
| Title | Title of the article | String | | |
| DOI | Digital object identifier of the article | String | | |
| # Exp | Experiment number | Positive integer | • The experiment number is an index to a specific experiment reported in a study, in case there are more than one. This index does not necessarily reflect the experiment number as reported in the paper, but does reflect the order of the experiments (i.e., if one of the experiments in the paper was not included in this database, the numbering would be different though the order will stay the same).<br>• When data from different samples were analyzed separately (e.g. studies using fMRI and EEG in different samples), they were encoded as separate experiments.<br>• Note that re-analyzed data collected in previous experiments was reported as an experiment. In case data from several experiments were combined and reanalyzed under the same framework, they were grouped and considered as one experiment. This was also the case when old data was compared with new one in the same analysis. For example, a paper reporting a new measure for predicting states of consciousness, based on a new analysis of data | |

| | | |
|---|---|---|
| | | collected in several older experiments (each using different techniques and populations), was encoded as a single experiment that used the techniques and populations of all re-analyzed experiments. |

*Exp. Paradigm*

| Main | The main paradigm/manipulation used in the experiment. A grouping variable that is used to denote different specific paradigms that belong to this group (the specific paradigm is coded in the next parameter in the table, Exp.Paradigm.Specific). | Abnormal Contents of Consciousness / Alcohol Consumption / Anesthesia / Attentional Manipulation / Caffeine Consumption / Case Study / Change Blindness / Cognitive Tasks / Competition (Binocular) / Competition (Monocular) / Computational Modelling / Contour Integration / Dichoptic Masking / Direct Stimulation / Disorders of Consciousness / Drowsiness / Emotion / Expectation / Familiarity / Family Nurture Intervention / Figure-Ground / Filling In / Fusion (Color) / Genetics / Illusions / Imagination / Lifespan / Locked In Syndrome / Masking / Mirror Masking / Motion Detection / Motion induced Blindness / N-Back / Pain / Passive Exposure / Pop out / Psychedelic Drugs / Repetition Suppression / Resting State / Reward / Sedation / Size Constancy / Sleep / Sperling Like / | • The value 'Case study' was used the classify experiments focusing on a specific individual, being compared to other individuals, or examined under a range of different conditions.<br>• The value 'Cognitive Tasks' refers to paradigms that are commonly used in cognitive research and are not specific to consciousness studies and do not directly manipulate consciousness (e.g., Mental rotation, Task switching, etc.).<br>• The 'Competition' paradigm was encoded as: Competition (Binocular) / Competition (Monocular) according to the specific paradigm used in the experiment.<br>• The value 'Direct Stimulation' refers to all paradigms in which brain activity was perturbed by the experimenters (e.g., using TMS).<br>• The value 'Sedation' (rather than 'Anesthesia') was used when subjects were not fully anesthetized but given anesthetic drugs at lower dozes so to achieve sedation.<br><br>Abbreviations: 'SWIFT' = semantic wavelet-induced frequency-tagging. | $Paradigm\_1 + \cdots + Paradigm\_n$<br>Where n denotes the number of different paradigms used in the experiment. So, if a specific experiment used both masking and CFS, it would be classified as masking + CFS |
|---|---|---|---|---|

| | | | | |
|---|---|---|---|---|
| | | Stimulus Degradation / SWIFT / Task Relevance / Visual Persistence / Visual Search | | |
| Specific | A sub-paradigm column, describing the type of the main paradigm. For example, for an experiment that uses "Masking" as the main paradigm, this column would indicate if it used Forward/Backward/Meta contrast/etc. masking. Note that in some cases only the main paradigm was coded (in cases where no specific paradigm was reported, or when the main paradigm that was used did not include sub paradigms, e.g., sleep). | (Organized according to main paradigms)<br>**Abnormal Contents of Consciousness:** Amusia / Blindsight / Conversion Deafness / Hypoactive delirium / Neglect / Parkinson Disease / Schizophrenia / Synesthesia / Tinnitus / Visual Extinction<br><br>**Anesthesia**: Dexmedetomidine / Dextromethorphan / Enflurane / Halothane / Isoflurane / Ketamine / Medetomidine / Midazolam / Pancuronium / Propofol / Sevoflurane / Sufentanil / Urethane / Xenon<br><br>**Attentional Manipulation**: Attentional Blink / Auditory Cueing / Crowding / Dichotic Listening / Distractor induced blindness / Dual Task / Inattentional Blindness / Selective Attention / Spatial Cueing<br><br>**Case Study**: Bilateral Frontal Affectation / Blindsight / Con version Deafness / Locked In Syndrome / Posterior Callosa l Section / Tinnitus / Unrespo | • Information about the main paradigm associated with each specific paradigm was provided in parenthesis along with additional information about the specific paradigm if relevant. Such additional information was given for the specific paradigms:<br>• *Attentional Blink* – the specific attentional blink paradigm used, if it differs from common paradigms (e.g., Attentional Blink (Attentional Manipulation, Ultra Rapid RSVP)).<br>• *Direct Stimulation* – a label describing the location in which the stimulation was given (e.g., TMS (Direct Stimulation, V1)). Note that the exact locations (e.g., in MNI coordinates) were not provided due to inconsistencies in the original reports of these locations.<br>• *Intensity Reduction* – when a specific intensity reduction method was reported (e.g., Intensity Reduction (Stimulus Degradation, Troxler).<br>• *Oddball* – the exact paradigm that was used – Local-Global/ Deviant detection/Visual Oddball / Roving Tactile Oddball/MMN.<br>• *Seizures* – a label describing the focal epileptic source if reported. As for the 'Direct Stimulation' value, a general label describing the epileptic source was provided rather than its exact location. | $SpecificParadigm_1(MainParadigm_j, AdditionalInformation_1 \,\&\, \ldots \,\&\, AdditionalInformation_k) + \cdots + SpecificParadigm_n (MainParadigm_m)$<br>Where n, m denotes the number of different specific paradigms and main paradigms used in the experiment.<br>The main paradigm associated with each specific paradigm is indicated in parenthesis, and the '&' sign is used to separate between different items of additional information, For Example, $TMS\ (DirectStimulation, V1 \,\&\, LOC) + IntensityReduction(Stimulus\ Degradation)$ |

| | | |
|---|---|---|
| | nsive Wakefulness Syndrome / Visual Extinction<br><br>**Cognitive Tasks**: Executive Control / Flankers Task / Language / Memory / Task Switching / Working Memory<br><br>**Competition**: b-CFS / Binocular Rivalry / Bistable percepts / CFS / Flash Supression<br><br>**Direct Stimulation**: Intracranial Stimulation / Olfactory Epithelium Stimulation / tACS / tDCS / TMS<br><br>**Disorders of Consciousness**: Coma / Emergence from MCS / Epilepsy / Minimal Consciousness State / Seizures /Unclassified DOC / Unresponsive Wakefulness Syndrome<br><br>**Expectation:** Emotional / Oddball / Prior Exposure<br><br>**Familiarity:** Own Name, Prior Exposure, Self-Face<br><br>**Illusions**: Apparent Motion / Benham's top Illusion / Flash-grab / Illusory Limb Movements / Illusory Motion Reversal / Illusory Rebound | • The 'Selective Attention' paradigm was used when a categorical non-spatial attentional cue was given to the participants.<br>• The 'Schizophrenia' and 'Parkinson Disease' values were grouped under the main paradigm 'Disorder of Consciousness' when they were discussed with relation to consciousness-related symptoms (e.g. hallucinations).<br>• The 'Unclassified DOC' value refers to experiments in which patients suffering from disorders of consciousness were examined, yet the specific disorders were not reported.<br><br>• Abbreviations: 'CFS' = continuous flash suppression, 'DOC' = disorders of consciousness, 'SAM' = ambiguous stroboscopic alternative motion, 'tACS' = transcranial alternating current stimulation, 'tDCS' = transcranial direct current stimulation, 'TMS' = transcranial magnetic stimulation. |

| | |
|---|---|
| | Motion / Implied Motion / SAM / Kanizsa / Moon Illusion / Octave Illusion / Pinna-Brelstaff / Proprioceptive Illusion / Tilt Illusion / Verbal Transformation / Zwicker Tone<br><br>**Imagination**: Motoric<br><br>**Masking**: Backward Masking / Dynamic Masking / Forward Masking / Metacontrast Masking / Object Substitution Masking / Paracontrast Masking / Target Masking<br><br>**Pop out:** Mooney Images<br><br>**Psychedelic Drugs**: Ketamine / Psilocybin<br><br>**Sedation**: Chloral Hydrate / Dexmedetomidine / Dextromethorphan / Halothane / Isoflurane / Ketamine / Lorazepam / Medetomidine / Midazolam / Nitrous Oxide / Propofol / Scopolamine / Urethane<br><br>**Stimulus Degradation**: Brief Presentation / Coherence Reduction / Intensity Reduction / Noise Induction<br><br>Abbreviations: 'CS' = conscious subjects (patients |

| | | | |
|---|---|---|---|
| | | used as controls in consciousness studies that do not have an active disorder of consciousness), 'UWS' = unresponsive wakeful subjects, 'MCS' = minimally conscious state, 'EMCS' = emergence from minimally conscious state, 'LIS' = locked in syndrome, 'PD' = Parkinson disease. | |
| Report/No-Report | Indicating whether a report or no-report paradigm was used | 0 (No-Report) / 1 (Report) / 2 (Report and No-Report) | • The value 'Report' was encoded for every experiment in which subjects were instructed to perform an active process of reporting or maintaining in memory information about the critical stimuli presented in the experiment (e.g., reporting the visibility of a stimulus, reporting confidence level in seeing a stimulus, or keeping count of the number of target stimuli presented during a block in the experiment).<br>• Experiments focusing on state consciousness were classified as 'No-Report' even if the state of consciousness was classified according to a common consciousness scale (e.g. SCR-R/ OAAS/ GCS/ other), sleep stages according to EEG or clinical examinations.<br>• 'Report and No-Report' value was classified when both report and no report conditions were included and analyzed separately in the same experiment. |

| | | | | |
|---|---|---|---|---|
| Content/State | Indicating whether the experiment focused on content / state consciousness or combined between the two | 0 (State) / 1 (Content) / 2 (State and Content) | • Experiments investigating Tinnitus/Synesthesia, searching for the correlates of the associated symptoms were classified as focusing on ’Content' consciousness.<br>• Experiments comparing stimuli processing under different states of consciousness (e.g., Local-Global experiments conducted on individuals with disorders of consciousness) were classified as focusing on 'State and Content'. | |
| *Sample* | | | | |
| Type | The type of population used in the experiment | 0 (Healthy Adults) / 1 (Healthy College Students) / 2 (Children) / 3 (Patients) / 5 (Nonhuman) / 6 (Computer) | • Experiments comparing patients and healthy subjects were classified with the value 'Patients' ('3'). Note that synesthetic subjects were classified as 'Healthy Adults' and not as 'Patients'.<br>• The value 'Computer' ('6') indicates that the experiment was based on computer simulations.<br>• Additional information about the population was given in parenthesis:<br>• *Children*/*Infants* – indicates the age cutoffs being used when those were explicitly reported.<br>• *Patients* – Information regarding the specific patient groups in the study was provided.<br>• *Non-Human* – The specific species studied. Computer simulations were not classified as 'Non-Human' (such experiments were classified as 'Computer').<br><br>Abbreviations: 'CS' = conscious subjects (patients used as controls in consciousness studies that do not have | $X\begin{pmatrix} AdditionalInformation_1 \dots \\ AdditionalInformation_n \end{pmatrix}$<br>Where $AdditionalInformation_i$ denotes additional information about population X for group i. For example:<br>3 (UWS Patients & MCS Patients & Healthy Controls) |

| | | | | |
|---|---|---|---|---|
| | | | an active disorder of consciousness), 'UWS' = unresponsive wakeful subjects, 'MCS' = minimally conscious state, 'EMCS' = emergence from minimally conscious state, 'LIS' = locked in syndrome, 'PD' = Parkinson disease.<br>• | |
| Total | Number of individuals that participated in the experiment | Positive Integer / NA | • When the sample was composed of subpopulations, additional information about their distribution was given in parenthesis. See the notes regarding the parameter Sample.Type for abbreviations.<br>• NA value indicates that the experiment was based on computer simulations or that the number of participants was not reported (occurred in a single experiment conducted on non-human animals). | $X\ (\#SubPopulation_1 + \cdots + \#SubPopulation_n)$<br>Where $\#SubPopulation_i$ denotes the number of subjects from subpopulation I.<br>For example: 20 (10 Blindsight Patients + 10 Healthy Controls). |
| Included | Number of included subjects in the experiment used for analysis (i.e., Sample size) | Positive Integer / NA | • NA value indicates that the experiment was based on computer simulations or that the number of participants was not reported (occurred in a single experiment conducted on non-human animals). | |

*Task*

| | | | | |
|---|---|---|---|---|
| Description | Verbal description of the task used in the experiment | String | | |
| Code | Numeral code for the type of task used in the experiment | 0 (No Task) / 1 (Discrimination) / 2 (Detection) / 3 (Go/No-Go) / 4 (Deviant Detection) / 5 (Stimulus Judgement) / 7 (Memory) / 8 (Change Detection) / 9 (Counting Task) / 10 (Delayed Task) / 14 (Task Switching) / 17 | • Computer simulations were classified as 'NA' (26), with the additional information: (Computer Simulations)<br>• Additional information about the task was given in parenthesis in specific cases:<br>• 'Memory' tasks, for which additional information include the following | $X_1\ (Additional\ Information_1) + \cdots + X_n\ (AdditionalInformation_n)$<br>Where $AdditionalInformation_i$ denotes additional information about the task $X_i$.<br>The + sign indicates that multiple tasks were used in the experiment.<br>For example:<br>22 (mixed percepts report included) |

| | | |
|---|---|---|
| | (NBack) / 18 (Dual Task) / 19 (Awareness Report) / 21 (Confidence Report) / 22 (Binocular Rivalry Task) / 23 (Visual Search) / 24 (Identification) / 26 (NA) / 27 (Driving) / 30 (Imagination) / 32 (Task Switching) / 33 (Mental Rotation) / 35 (Color Matching) / 36 (Mood) / 37 (Mathematics) | values: Episodic, Old/New, Recall and Match to Sample.<br>• 'Identification' tasks, for which additional information include the following values: Naming, Silent Naming, Match to Sample, Recall.<br>• Note that 'Match to Sample' and 'Recall' tasks were classified as specific tasks under 'Memory' and 'Identification' codes. The value 'Memory' was used only when the participants were asked to memorize the target stimuli. Otherwise (e.g., when subjects were asked to identify which of two stimuli was presented in the trial), the task was encoded as 'Identification'.<br>• The value 'Binocular Rivalry Task' (22) was complemented with the additional information: 'Mixed percept report' when mixed percept report was included.<br>• The value 'Stimulus Judgement' (5) was complemented with additional information regarding the specific stimulus type. |

### *Stimuli*

| | | | | |
|---|---|---|---|---|
| Description | A verbal description of the stimuli that were used in the experiment | | | |
| Category | Category of the stimuli that were used in the experiment. | Animals / Artificial Scenes / Bodies / Checkerboard / Chinese Pictographs / Color / Contours / Digits / Drawings / Electric Stimulation / Epithelium Stimulation / Faces / Figure-Ground / Geometric Shapes / Gratings | • Note that the encoded stimulus category refers to the main stimuli presented in the experiment. In addition to the critical stimuli presented in the experiment, additional stimuli categories were added according to their relevance (e.g., experiments using face stimuli | $X_1\ (Additional\ Information_1) + \cdots + X_n\ (AdditionalInformation_n)$<br>Where $AdditionalInformation_i$ denotes additional information about the stimuli category $X_i$.<br>The + sign indicates that stimuli from multiple categories were used in the experiment.<br>For example: |

| | | | | |
|---|---|---|---|---|
| | | / Kanizsa / Landolt / Letters / Light Flashes / Motion / Music / Natural Scenes / Nociceptive stimulation / Noise / None / Numbers / Objects / Pacman / Patterns / Pneumatic stimulations / Real Objects / Sexual Images / Sounds / Speech / Symbols / Textures / Verniers / Videos / Virtual Reality Objects / Words | as their critical stimuli were categorized as 'Faces + Real Scenes' if the faces were presented within real scenes. Similarly, experiments using face stimuli as targets and houses as distractors were coded as "Faces + Houses").<br>• Additional information about the specific stimuli type that was provided for geometric shapes (Arrows, Motion, Lines, Disc, Dots, Rings, Bars Circles, Rectangles, Squares, Diamonds) and objects (Houses, Clocks, Domino) and virtual reality objects (Houses, Road, Moon).<br>• The value 'Colors' indicates that the main feature of the stimuli in the experiment was color. Accordingly, colorful and grayscale stimuli were not classified differently.<br>• The value 'None' indicates that no stimuli were used in the experiment. | Geometric Shapes (Lines) + Words |
| Modality | Modality in which the stimuli were presented in the experiment | Auditory / None / Olfactory / Tactile / Visual | • Note that this parameter relates only to the stimuli being used by the researchers, and not to the resulting perceptual states (e.g., experiments using tactile stimulation to induce proprioceptive illusions were classified in this parameter as 'Tactile' and not 'Proprioception').<br>• The value 'None' indicates that no stimuli were used in the experiment. | $X_1 + \cdots + X_n$<br>Where $X_i$ denotes a specific stimulus modality used in the experiment and + indicates that multiple stimuli modalities were used. |
| Duration | Duration of critical stimuli presentation | Time in ms / NA / String / None | • Mapping between the duration of the stimuli and the modality of the category was given in parenthesis for cross-modal experiments<br>• A string code was used when stimuli were presented in multiple durations | $X_1\ (Modality_j) + \cdots + X_n\ (Modality_m)$<br><br>Where $Modality_j$ denotes the modality of the stimuli that were presented for $X_i$ duration. |

| | | | | |
|---|---|---|---|---|
| | | | or when additional information was needed to disentangle complex duration reports.<br>• The value 'NA' indicates that the duration of the stimuli presentation was not reported.<br>• The value 'None' indicates that no stimuli were used in the experiment. | |
| Contrast | Contrast of the stimuli that were used in the experiment | Average contrast / NA / String / None | • Note that when contrast was not reported as a value, or when it varied between subjects (e.g., when contrast was manipulated to achieve 70% performance), a verbal description was given, rather than a numeral one.<br>• In auditory experiments, the contrast was reported using a dB scale.<br>• The value 'None' indicates that no stimuli were used in the experiment.<br>• The value 'NA' indicates that the contrast of the stimuli presentation was not reported. | $X_1\ (Modality_j) + \cdots + X_n\ (Modality_m)$<br><br>Where $Modality_j$ denotes the modality of the stimuli that were presented at a contrast of $X_i$. |

### *Consciousness Measures*

| | | | | |
|---|---|---|---|---|
| Description | Verbal description of the consciousness measure used in the experiment. Note that in studies focusing on state consciousness, this column included information about the state classification procedure (usually according to a common scale) | String | | |
| Type | Timing of consciousness measure used in the experiment. | None / Post Experiment / Pre Experiment / Separate Experiment / Trial By Trial | • A 'None' value indicates that no consciousness measure was taken during the experiment. | $X_1 + \cdots + X_n$<br>Where $X_i$ denotes a specific type of measure of consciousness used in the experiment and + |

| | | | | |
|---|---|---|---|---|
| | | | • The value 'Separate Experiment' indicates that awareness was measured on a separate sample than the sample analyzed in the main experiment. | indicates that multiple consciousness measures of different types were used. |
| Taken | Type of consciousness measure | Condition Assessment / None / Objective / Sleep Monitoring / State induction Assessment / Subjective | • Experiments in which measures were used to categorize a consciousness relevant participants' condition (e.g., categorization of patients to disorders of consciousness) were classified as using a 'Condition Assessment' measure.<br>• Experiments that adjusted anesthetic drug dosages until reaching a target consciousness state were classified as 'State Induction Assessment'.<br>• Experiments in which the measure of consciousness was described in terms of confidence were classified as 'Subjective (Confidence)'.<br>• Experiments measuring consciousness subjectively yet also using catch trials (trials in which the critical stimulus was absent) to conduct signal detection theory (SDT) analysis of awareness were classified as 'Objective and Subjective' | |
| *Techniques* | Neuroscientific techniques used in the experiment | Ca2 Imaging / Computational Modelling / EEG / fMRI / Intracranial EEG / Intracranial Stimulation / MEG / MRI / PET / tACS / tDCS / TMS | • Experiments using several neuroscientific techniques were classified as such, even when only one technique was used to measure NCCs (e.g., experiments using both fMRI and EEG, where EEG is used to classify sleep stages and fMRI is used to measure the dependent variable). | |

| | | | |
|---|---|---|---|
| | | | Abbreviations: 'Ca2 Imaging' = calcium imaging, 'EEG' = electroencephalography, 'TMS' = transcranial magnetic stimulation, 'fMRI' = functional magnetic resonance imaging, 'MRI' = magnetic resonance imaging, 'MEG' = magnetoencephalography, 'PET' = positron emission tomography, 'tACS' = transcranial alternating current stimulation, 'tDCS' = transcranial direct current stimulation |

### *Findings*

| | | | | |
|---|---|---|---|---|
| Summary | Verbal description of the findings of the experiment. Complements the NCC Tags column and describes the main findings at more length. | String | | |
| NCC Tags | Numeral codes for specific NCC candidates. | 0 (Frontal) / 1 (Ventral Stream) / 2 (V1) / 3 (P300) / 4 (VAN) / 5 (Gamma) / 6 (Complexity) / 7 (Local Synchronization) / 8 (Global Synchronization) / 9 (Fronto Parietal connectivity) / 10 (Variability) / 11 (A1) / 12 (Dorsal Stream) / 13 (Beta) / 14 (Alpha) / 15 (CNV) / 16 (Parietal) / 17 (DMN) / 18 (Small Worldness) / 19 (PHI Approximation) / 20 (Metacognition) / 21 (Posterior) / 22 (N2pc) / 23 (Recurrent Processing) / 24 (GABA/NMDA) / 25 (P2) / 26 (MMN) / 27 (N2) / 28 | • Generally, the classification of findings was based on the original reports of the authors. No attempts were made to assess the reliability or the validity of the reported findings in both conceptual and statistical manner.<br>• Additional information about each finding was given in parenthesis. This information includes different values according to the specific findings reported (see the 'Encoding format' for details).<br>• Note that the spatial and temporal labels used to encode this parameter also reflect the way the results were described by the authors. We did not | $X_1 \begin{pmatrix} AdditionalInformation_1 \\ AdditionalInformation_2 \end{pmatrix} + X_n \ (AdditionalInformation_n)$<br><br>Where $X_i$ denotes a specific finding of the experiment and + indicates that multiple findings were found. The sign – before any $X$ indicates that this finding was interpreted as 'negative evidence' (see Notes). The sign & indicates multiple findings under the same NCC tag code. Note that the additional information provided for each $X$ is different.<br><br>For EEG component findings:<br>$TimeWindow_i ms$<br>$< Technique_i >$<br>Where $TimeWindow_i$ is encoded with the format $onset - offset\ ms$ to indicate the exact time |

| | | | |
|---|---|---|---|
| | (Theta) / 29 (Delta) / 30 (ART) / 31 (V4) / 32 (Early Components) / 33 (Late Components) / 34 (Temporal Parietal Connectivity) / 35 (S1) / 36 (N140) / 37 (N170) / 38 (Centrality) / 39 (N1) / 40 (CFC) / 41 (Anterior Posterior Connectivity) / 42 (Subcortical structures) / 43 (Cortical Subcortical connectivity) / 44 (Acetylcholine) / 46 (SN) / 47 (Motor areas connectivity) / 48 (Temporal Occipital connectivity) / 49 (Prestimulus Components) / 50 (Low frequencies <1Hz) / 51 (Uncinate Fasciculus) / 53 (SPCN) / 55 (Figure Ground Difference) / 56 (Border Difference) / 57 (P1) / 58 (Frequency Increase) / 59 (Sleep Spindles) / 60 (Slow Waves Activity) / 62 (ERN) / 63 (EPN) / 64 (Hyper Synchronization) / 65 (Plasticity) / 66 (Anatomic Functional connectivity similarity) / 67 (Ultra slow fluctuations) / 69 (M70) / 70 (M130) / 71 (ARN) / 72 (M280) / 74 (N400) / 75 (Pe) / 76 (Change related positivity) / 77 (N150) | interpret or assessed the validity of the original report.<br>• The different values used for spatial findings (e.g., 'Frontal', 'Parietal', 'Posterior', etc.) were used as grouping variables and were not used for analysis. Only the encoded AAL3 label and coordinates were considered as the markers of spatial findings. See column 'Findings.Spatial AAL Mapping' for the encoding of specific AAL coordinates for fMRI findings.<br>• Note that some findings were classified as a 'negative finding'. 'Negative finding' indicates that the authors reported *not* finding a certain candidate NCC in their data, or finding evidence against a certain candidate NCC (e.g., finding a specific NCC candidate for unconscious trials). Findings that were only interpreted as indexing other processes and not consciousness per se, without experimentally dissociating these processes from conscious processing were not classified as 'Negative findings' (e.g., significant P300 found in the contrast between seen and unseen trials which were interpreted in the discussion as related to post-perceptual processes, yet without having any manipulation in the experiment providing evidence that this is indeed the case was classified as 'P300', and not as 'negative P300').<br>• Marginally significant findings were classified according to the | window associated with the finding. When a single time point was reported (for example for peak amplitude measures), the offset was omitted. An optional code: $Technique_i$ was used to encode the mapping to a specific technique in experiments that used multiple techniques.<br><br>For Frequency findings:<br>$Type_i\ Band_iHz\ Correlation\ Sign_i < TimeWindow_ims > < Technique_j >$<br>Where $Type_i \in \{Power, Connectivity, Phi, Complexity, TE, PCA, LRTC, Microstates, CD, Clustering\}$ (TE = transfer entropy, PCA = principal components analysis, LRTC = long-range temporal correlations, CD = correlation dimension) and indicates that type of frequency finding reported in the experiment. $Band_i$ is encoded with the format $lowerBandFrequency - HigherBandFrequencyms$ to indicate the exact band associated with the finding. When a specific frequency was reported the higher band frequency was omitted. $Sign_i$ indicates whether the frequency finding is negatively or positively correlated with consciousness. $TimeWindow_i$ is optional and was encoded with the format $onset - offset\ ms$ to indicate the exact time window associated with the finding. When a single time point was reported (for example for peak amplitude measures), the offset was omitted. An optional code: $Technique_j$ was used to encode the mapping to a specific technique in experiments that used multiple techniques.<br><br>For spatial fMRI findings:<br>$AAL3Label_i < Technique_j >$ |

| | | | |
|---|---|---|---|
| | | interpretation of the authors (i.e., if the authors reported them as marginal and considered them as a finding, also referring to them in the discussion etc., they were classified as a finding). Such cases are marked with the 'Marginal' label in the respective Findings.Notes column.<br>• Connectivity findings were grouped according to the broad spatial properties of the reported finding. For example, the value 'Fronto-Parietal Connectivity' was used in experiments reporting breaking of connectivity between prefrontal and parietal areas in the transition between conscious and unconscious states, whereas the value 'Anterior–Posterior Connectivity' was used when such connectivity correlate was found between temporal/occipital and frontal areas.<br>• The value 'Centrality' codes for findings related to the graph-theoretical measures of centrality (betweenness, eigenvector, degree, etc.).<br>• The values 'Figure-Ground difference' and ' Boarder difference' encode specific EEG components which were reported<br>• Frequency findings were encoded according to the frequency band label reported by the authors (see the additional information for the specific frequency band that was analyzed). When no such band label was reported by the authors, the band label | Where $AAL3Label_i$ indicates the AAL3 atlas label associated with the findings. The label was encoded based on the MNI coordinates reported for each finding (for results reported in other coordinate systems e.g., Talairach, the coordinates were transformed to MNI space).<br>An optional code: $Technique_j$ was used to encode the mapping to a specific technique in experiments that used multiple techniques.<br><br>For other spatial findings:<br>$Label\ \#\ comment\ including\ electrodes$<br>Spatial findings that were found in non-fMRI studies were encoded with less accuracy. When source localization or direct stimulation were used, the area label reported in the study was encoded, in other cases, an attempt was made to provide any informative label.<br>Where Label<br><br>Other findings:<br>$Free\ text$<br>For other findings, the additional information could include free text which was not analyzed in this review. |

| | | |
|---|---|---|
| | | was classified according to the common standard in the field.<br>• The values 'GABA/NMDA' and 'Acetylcholine' were encoded when they were directly found and reported as NCC candidates.<br>• The values 'Global Synchronization' and 'Local Synchronization' indicate that the authors interpreted their findings as indicating an effect of global/local, long-range/local, activity, respectively. For example, experiments using the Local-Global paradigm, and findings a global effect to be correlated with consciousness were encoded with the 'Global Synchronization' finding, while other experiments finding network modularity to be correlated with consciousness were encoded with a 'Local Synchronization' finding.<br>• The values 'Late Components', 'Early Components' and 'Prestimulus Components' were used when a temporal component was found to be correlated with consciousness, yet no specific component was reported by the authors. In these cases, the component was classified as Early/ Late according to the interpretation of the authors. For example, in experiments using TMS at separate time windows, and finding that consciousness is modulated by perturbation on specific time windows (late, early, or before stimulus presentation), the finding was encoded according to the temporal |

| | | | | |
|---|---|---|---|---|
| | | | properties of the time window, according to the way the authors classified the finding in their paper (e.g., reflecting late/early processes).<br>• The 'P300' value was used for P3a and P3b components, as well as in cases in which researchers referred to MEG components as stemming from activity in the generators P300.<br>• The value 'Recurrent Processing' was used when the authors explicitly interpreted their findings as reflecting recurrent processing. For example, in experiments using TMS when the researchers perturb neuronal activity at time windows associated with recurrent processing, and finding resulting significant modulation of consciousness, the findings were classified as 'Recurrent Processing'.<br><br>Abbreviations: 'VAN' = visual awareness negativity, 'CNV' = contingent negative variation, 'CFC' = cross frequency correlation, 'SN' = selection negativity, 'SPCN' = sustained posterior contralateral negativity, 'ERN' = error related negativity, 'ARN' = auditory awareness related negativity. | |
| Measures | Numeral codes for the measures taken in the experiment to analyze NCC findings. | 0 (Decoding) / 1 (BOLD) / 2 (Frequencies) / 3 (ERP) / 4 (Mutual Information) / 5 (Synchronization) / 6 (Behavioral (Accuracy)) / 7 (Behavioral (RT)) / 9 (Connectivity) / 10 (PHI) / 11 (Graph theoretical measures) / 14 (Entropy) / 15 (Global | • Additional information about the measure was given in parenthesis.<br>• The value 'Connectivity' groups together different kinds of measures and analyses. Additional information specifies whether effective / functional/ structural connectivity was measured. Additional information is also provided regarding the specific | $X_1\ (AdditionalInformatiom_1) + X_n\ (AdditionalInformation_n)$<br><br>Where $X_i$ denotes a specific measure used in the experiment and + indicates that multiple measures were used. The sign & indicates multiple findings under the same NCC tag code. |

| | | | |
|---|---|---|---|
| | Field Power) / 16 (PCA) / 17 (Lempel Ziv) / 18 (H2_15O) / 19 (Variability) / 20 (Adaptation) / 21 (Metacognition) / 22 (Visibility) / 25 (Dimension of activation) / 26 (fALFF) / 27 (18F Fluorodeoxyglucose) / 28 (CFC) / 29 (LRTC) / 30 (Calcium Imaging) / 31 (K Complex) / 32 (TCT) / 33 (DISS) / 34 (Observation) / 35 (Microstates) / 36 (Stimulation Reactivity) / 37 (Frequency Tagging) / 39 (Hopf bifurcation parameter) / 41 (Slow Wave Activity) / 42 (Auto Information Flow) / 43 (Cross Information Flow ciF) / 44 (Auto Correlation) / 45 (Topo) / 46 (PCI) / 47 (Physiological Measure) / 49 (Computer Simulations) / 50 (Phosphene Threshold) / 51 (Frequency Change Index) / 52 (Brain Behavior Correlation) / 56 (Nonlinear correlation index) / 57 (DSI) / 58 (Complexity of functional connectivity) / 63 (BIS) / 64 (Correlation dimension) / 65 (Dominance) / 66 (Epileptogenicity Index) / 67 (Spike Suppression) / 68 (Mean Dwell Time) / 69 (Network Backbones) | analysis used (e.g., phase lag index (PLI), psychophysiological interaction (PPI) / fractional anisotropy (FA)/ etc.).<br>• The value 'Decoding' was used to indicate that brain activity was used to decode a certain variable in the experiment (e.g., the consciousness state or conscious report of a subject or alternatively an experimentally manipulated variable such as stimuli category). Additional information provides details about the specific analysis that was conducted (e.g., MVPA\ SVM \ LDA\ etc.).<br>• For the value 'ERP' the additional information indicates whether the analysis was performed according to the peak (e.g., comparing peak amplitude/latency), within a time window (e.g., mean amplitude analysis) or according to other methods (e.g., analyzing the latency of 50% peak amplitude/ etc.). When sample by sample analysis was conducted the entire significant time window was classified, and the tag 'Cluster' was added.<br>• For the value 'Frequencies' the additional information indicates whether induced/evoked/resting state based analysis was used.<br>• The value 'Observation' indicates that the researchers observed subjects' behavior to make inferences about their consciousness state. For example, experiments in which the symptoms of patients were monitored | Note that the additional information provided for each $X$ is different, yet the details provided for each classification were not used for analysis.<br>For example:<br>0 (MVPA) + 9 (functional connectivity) + 1 |

| | | | | |
|---|---|---|---|---|
| | | | as an index of consciousness disorders status, were classified as 'Observation'.<br>• Note that the values 'Synchronization' and 'Connectivity' were frequently classified together, relating to the same measure. For example, the measure phase lag index (PLI) which measures phase synchronization of two signals is commonly used as a measure of functional connectivity. As such, experiments using PLI to measure functional connectivity were classified as 'Synchronization (PLI) + Connectivity (functional connectivity, PLI).<br><br>Abbreviations: 'BOLD' = blood oxygen level dependent, 'ERP' = event related potentials, 'RT = reaction time, 'DTI-FA' = diffusion tensor imaging – fractional anisotropy, 'PCA'= principal components analysis, 'DCM' = dynamic causal modeling, 'fALFF' = fractional amplitude of low-frequency fluctuations, 'CFC' = cross frequency correlation, 'LRTC' = long-range temporal correlation, 'TCT' = topographic consistency test , 'DISS' = global dissimilarity index, 'Topo' = topographic similarity , 'PCI' = perturbational complexity index ,'DSI' = diffusion spectrum imaging ,'BIS' = Bispectral index. | |
| Spatial AAL Mapping | Information about spatial fMRI findings. Specifically, the column encodes the AAL3 atlas | See the format column for the formula to generate valid values. | See the description of the column 'Findings.NCC Tags' for details about the 'Label' code. | $AAL3Label_1 < Label_1\ X_1, Y_1, Z_1 \% \ldots \% X_m, Y_m, Z_m > + AAL3Label_n < Label\ X, Y, Z >$ |

| | | | | |
|---|---|---|---|---|
| | label and respective MNI coordinates for each finding in the experiment | | | Where $AAL3Label_i$ denotes an AAL3 atlas label of a brain area reported in an fMRI study and + indicates that multiple brain areas were found in this experiment. $Label_i$ indicates the label used to encode the finding in the 'Findings.NCC Tags' column and the exact AAL3 atlas coordinates that were mapped to the reported fMRI results. The sign & indicates that multiple brain activations were found for a similar label. |
| Encoding Notes | Verbal notes about the classification. Includes information about non-intuitive aspects of the classification process / the interpretations of the findings. | String | | X |
| *Interpretation* | | | | |
| GNW | A classification of the experiment as supporting/challenging GNW. | 0 (Against) / X (Neutral) / 1 (Pro) | Here and below, support/challenge was classified solely based on the way the authors described their findings in the discussion (that is, we did not engage in interpretation of findings). | |
| HOT | A classification of the experiment as supporting/challenging HOT. | | | |
| IIT | A classification of the experiment as supporting/challenging IIT. | | | |
| RPT | A classification of the experiment as supporting/challenging RPT. | | | |
| *Theory Driven* | An indicator of whether the experiment was presented as testing one or more of the theories. | 0 (Post hoc) / 1 (Mentioned the theories) / 2 (Theory driven) | • The 'Theory driven' value indicates that the experiment was a-priori designed to test the predictions of at least one theory. This is inferred from | $X\ (Theory_1\ \&\ \ldots\ \&\ Theory_n)$ Where $X$ is a single value, being one of the theory driven codes (0/1/2). The theories t0 which X and the sign & indicates multiple theories that |

| | | | | |
|---|---|---|---|---|
| | | | the introduction: if the authors present the study in the introduction as testing a prediction by at least one theory, it would be classified as ‘Theory driven’.<br>• The 'Post hoc' value indicates that the results of the experiment were only interpreted post hoc in the discussion.<br>• The value 'Mentioned the theories' indicates that the experiment was not theory-driven, yet the authors mentioned one of the theories in the introduction. | were mentioned (for code '1' – mentioning a theory) or the theories that were investigated in the experiment (for code '2' – theory driven). |
| *Internal Replication* | An indicator for whether the experiment is an internal replication of a previous experiment reported in the same article | 0 (Not an internal replication) / 1 (Internal replication) | | |
| *Metadata* | | | | |
| Title | Title of the article | String | All the *Metadata* parameters were extracted automatically from the Scopus database via the Scopus website interface. | |
| DOI | Digital object identifier of the article | String | | |
| Authors | Author names | String | | |
| Publication Year | Publication year | Integer | | |
| Link | URL link to the Scopus page of the article | URL | | |
| Source Title | Name of the journal in which the article was published | String | | |
| Cited By | How many papers cited the article according to Scopus' count | Integer | | |

| | | |
|---|---|---|
| Abstract | Full abstract | String |
| Affiliations | Author affiliations | String |
| Author Keywords | Keywords as given by the authors | String |
| Index Keywords | Keywords as given by Scopus | String |
| Funding Details | Names of the foundations/institutions that funded the study | String |
| References | List of the article's references | String |
| Publisher | Name of the publisher of the article | String |
| Abbreviated Source Title | Abbreviated of the journal name | String |

*Table S4*. A full description of the extraction sheet was used for the systematic review.

**Supplementary Table S5: Feature importance – Main analysis after exclusion**

| <u>Feature</u> | <u>5%</u> | <u>95%</u> | <u>Mean Importance</u> |
|---|---|---|---|
| State (State vs. Content) | 0.061 | 0.085 | 0.073 |
| No-Report (Report vs. No-Report) | 0.036 | 0.058 | 0.046 |
| Connectivity (Dependent Measures) | 0.028 | 0.046 | 0.037 |
| Subjective (Consciousness Measures Type) | 0.027 | 0.042 | 0.034 |
| Patients (Population) | 0.017 | 0.031 | 0.024 |
| Frequencies (Dependent Measures) | 0.015 | 0.028 | 0.021 |
| Random Variable | 0.012 | 0.028 | 0.019 |
| TMS (Neuroscientific Techniques) | 0.009 | 0.021 | 0.015 |
| Geometric Shapes (Stimuli Categories) | 0.010 | 0.019 | 0.015 |
| Entropy (Dependent Measures) | 0.009 | 0.017 | 0.013 |
| ERP (Dependent Measures) | 0.008 | 0.017 | 0.013 |
| Discrimination (Task) | 0.004 | 0.013 | 0.008 |
| fMRI (Neuroscientific Techniques) | 0.003 | 0.013 | 0.008 |
| Propofol (Specific Paradigm) | 0.004 | 0.011 | 0.007 |
| Intracranial EEG (Neuroscientific Techniques) | 0.004 | 0.011 | 0.007 |
| Intensity Reduction (Specific Paradigm) | 0.003 | 0.011 | 0.007 |
| Detection (Task) | 0.003 | 0.010 | 0.006 |
| Synchronization (Dependent Measures) | 0.002 | 0.009 | 0.005 |
| Metacontrast Masking (Specific Paradigm) | 0.004 | 0.007 | 0.005 |
| Auditory (Stimuli Modality) | 0.002 | 0.009 | 0.005 |
| Brain Behavior Correlation (Dependent Measures) | 0.002 | 0.007 | 0.005 |
| Color (Stimuli Categories) | 0.003 | 0.007 | 0.005 |
| Metacognition (Dependent Measures) | 0.003 | 0.006 | 0.004 |
| Isoflurane (Specific Paradigm) | 0.002 | 0.006 | 0.004 |
| Faces (Stimuli Categories) | 0.002 | 0.006 | 0.004 |
| Graph theoretical measures (Dependent Measures) | 0.002 | 0.006 | 0.004 |
| Words (Stimuli Categories) | 0.002 | 0.006 | 0.004 |
| Global Field Power (Dependent Measures) | 0.002 | 0.005 | 0.003 |
| Digits (Stimuli Categories) | 0.001 | 0.006 | 0.003 |
| Decoding (Dependent Measures) | 0.002 | 0.005 | 0.003 |
| Stimulation Reactivity (Dependent Measures) | 0.001 | 0.006 | 0.003 |
| Oddball (Specific Paradigm) | 0.001 | 0.006 | 0.003 |

| | | | |
|---|---|---|---|
| Backward Masking (Specific Paradigm) | 0.001 | 0.005 | 0.003 |
| Gratings (Stimuli Categories) | 0.001 | 0.004 | 0.003 |
| Brief Presentation (Specific Paradigm) | 0.001 | 0.004 | 0.002 |
| MEG (Neuroscientific Techniques) | 0.001 | 0.004 | 0.002 |
| Mutual Information (Dependent Measures) | 0.001 | 0.003 | 0.002 |
| Emotion (Main Paradigm) | 0.001 | 0.003 | 0.002 |
| Memory (Task) | 0.001 | 0.003 | 0.002 |
| Inattentional Blindness (Specific Paradigm) | 0.001 | 0.003 | 0.002 |
| Non-human (Population) | 0.000 | 0.004 | 0.002 |
| Animals (Stimuli Categories) | 0.000 | 0.003 | 0.002 |
| Delayed Task (Task) | 0.001 | 0.002 | 0.002 |
| Change Detection (Task) | 0.001 | 0.002 | 0.001 |
| Counting task (Task) | 0.001 | 0.002 | 0.001 |
| Letters (Stimuli Categories) | 0.000 | 0.002 | 0.001 |
| Tactile (Stimuli Modality) | 0.000 | 0.002 | 0.001 |
| Noise Induction (Specific Paradigm) | 0.000 | 0.002 | 0.001 |
| Natural Scenes (Stimuli Categories) | 0.000 | 0.002 | 0.001 |
| Checkerboard (Stimuli Categories) | 0.001 | 0.001 | 0.001 |
| Awareness Report (Task) | 0.000 | 0.002 | 0.001 |
| Objects (Stimuli Categories) | 0.000 | 0.002 | 0.001 |
| Binocular Rivalry (Specific Paradigm) | 0.000 | 0.002 | 0.001 |
| Attentional Blink (Specific Paradigm) | 0.000 | 0.002 | 0.000 |
| Identification (Task) | 0.000 | 0.001 | 0.000 |
| PHI (Dependent Measures) | 0.000 | 0.002 | 0.000 |
| Stimulus Judgement (Task) | 0.000 | 0.001 | 0.000 |
| Passive Exposure (Main Paradigm) | 0.000 | 0.001 | 0.000 |
| Forward Masking (Specific Paradigm) | 0.000 | 0.001 | 0.000 |

*Table S5*. Importance factors calculated for each parameter used to train the random forest classifier, trained to classify theory-support according to methodological parameters extracted from each experiment in the database (see Figure 4a-b in the main article). Importance values indicate hamming loss importance scores calculated using a permutation method. The permutation method was repeated 1000 times creating a distribution of importance scores per feature. The columns '5%' and '95%' and 'Mean Importance' denote the $5^{th}$, $95^{th,}$ and the mean importance scores in the distribution of each feature.

**Supplementary Table S6: Feature importance – full model**

| Feature | 5% | 95% | Mean Importance |
|---|---|---|---|
| Subjective (Consciousness Measures Type) | 0.015 | 0.029 | 0.022 |
| Frequencies (Dependent Measures) | 0.010 | 0.021 | 0.015 |
| Connectivity (Dependent Measures) | 0.007 | 0.018 | 0.012 |
| Random Variable | 0.006 | 0.018 | 0.012 |
| State (State vs. Content) | 0.003 | 0.013 | 0.008 |
| Geometric Shapes (Stimuli Categories) | 0.003 | 0.011 | 0.007 |
| Resting State (Main Paradigm)* | 0.002 | 0.009 | 0.005 |
| Entropy (Dependent Measures) | 0.002 | 0.008 | 0.005 |
| Intensity Reduction (Specific Paradigm) | 0.002 | 0.007 | 0.005 |
| Behavioral (Accuracy) (Dependent Measures)* | 0.002 | 0.006 | 0.004 |
| Discrimination (Task) | 0.002 | 0.006 | 0.004 |
| Synchronization (Dependent Measures) | 0.002 | 0.006 | 0.004 |
| Visual (Stimuli Modality)* | 0.001 | 0.006 | 0.004 |
| Objective (Consciousness Measures Type)* | 0.001 | 0.007 | 0.004 |
| Content (State vs. Content)* | 0.001 | 0.007 | 0.004 |
| Color (Stimuli Categories) | 0.002 | 0.005 | 0.004 |
| Detection (Task) | 0.002 | 0.006 | 0.003 |
| Words (Stimuli Categories) | 0.002 | 0.006 | 0.003 |
| Metacognition (Dependent Measures) | 0.002 | 0.005 | 0.003 |
| Patients (Population) | 0.001 | 0.006 | 0.003 |
| Metacontrast Masking (Specific Paradigm) | 0.002 | 0.004 | 0.003 |
| EEG (Neuroscientific Techniques)* | 0.001 | 0.006 | 0.003 |
| TMS (Specific Paradigm)* | 0.001 | 0.005 | 0.003 |
| Brain Behavior Correlation (Dependent Measures)* | 0.002 | 0.004 | 0.003 |
| Propofol (Specific Paradigm) | 0.001 | 0.005 | 0.003 |
| Behavioral (RT) (Dependent Measures) | 0.002 | 0.003 | 0.002 |
| Decoding (Dependent Measures) | 0.001 | 0.004 | 0.002 |
| ERP (Dependent Measures) | 0.001 | 0.004 | 0.002 |
| Graph theoretical measures (Dependent Measures) | 0.000 | 0.004 | 0.002 |
| Healthy Adults (Population)* | 0.000 | 0.005 | 0.002 |
| Digits (Stimuli Categories) | 0.000 | 0.004 | 0.002 |

| | | | |
|---|---|---|---|
| Gratings (Stimuli Categories) | 0.001 | 0.003 | 0.002 |
| Auditory (Stimuli Modality)* | 0.000 | 0.005 | 0.002 |
| Minimal Consciousness State (Specific Paradigm)* | 0.000 | 0.003 | 0.002 |
| Delayed Task (Task) | 0.001 | 0.002 | 0.002 |
| Isoflurane (Specific Paradigm) | 0.001 | 0.002 | 0.002 |
| Faces (Stimuli Categories) | 0.001 | 0.003 | 0.002 |
| Pre Experiment (Consciousness Measures Phase)* | 0.001 | 0.002 | 0.002 |
| Inattentional Blindness (Specific Paradigm) | 0.001 | 0.002 | 0.002 |
| Animals (Stimuli Categories) | 0.000 | 0.003 | 0.001 |
| Global Field Power (Dependent Measures) | 0.000 | 0.003 | 0.001 |
| Backward Masking (Specific Paradigm) | 0.000 | 0.003 | 0.001 |
| TMS (Neuroscientific Techniques) | 0.000 | 0.003 | 0.001 |
| Brief Presentation (Specific Paradigm) | 0.000 | 0.002 | 0.001 |
| Counting task (Task) | 0.001 | 0.002 | 0.001 |
| Memory (Task) | 0.001 | 0.002 | 0.001 |
| Emotion (Main Paradigm) | 0.001 | 0.002 | 0.001 |
| Noise Induction (Specific Paradigm) | 0.001 | 0.002 | 0.001 |
| No-Report (Report vs. No-Report) | 0.000 | 0.002 | 0.001 |
| Non-human (Population) | 0.000 | 0.002 | 0.001 |
| BOLD (Dependent Measures)* | 0.000 | 0.002 | 0.001 |
| Mutual Information (Dependent Measures) | 0.000 | 0.002 | 0.001 |
| Natural Scenes (Stimuli Categories) | 0.000 | 0.002 | 0.001 |
| Change Detection (Task) | 0.000 | 0.002 | 0.001 |
| Stimulation Reactivity (Dependent Measures) | 0.000 | 0.002 | 0.001 |
| Post Experiment (Consciousness Measures Phase)* | 0.000 | 0.002 | 0.001 |
| No Task (Task)* | 0.000 | 0.002 | 0.001 |
| Checkerboard (Stimuli Categories) | 0.001 | 0.001 | 0.001 |
| Intracranial EEG (Neuroscientific Techniques) | 0.000 | 0.002 | 0.001 |
| MEG (Neuroscientific Techniques) | 0.000 | 0.002 | 0.001 |
| Report (Report vs. No-Report)* | -0.001 | 0.002 | 0.001 |
| Awareness Report (Task) | 0.000 | 0.001 | 0.001 |
| Identification (Task) | 0.000 | 0.002 | 0.001 |
| fMRI (Neuroscientific Techniques) | 0.000 | 0.002 | 0.001 |
| Objects (Stimuli Categories) | 0.000 | 0.002 | 0.000 |
| Binocular Rivalry (Specific Paradigm) | 0.000 | 0.001 | 0.000 |

| | | | |
|---|---|---|---|
| None (Stimuli Modality)* | -0.001 | 0.002 | 0.000 |
| Stimulus Judgement (Task) | 0.000 | 0.001 | 0.000 |
| Trial By Trial (Consciousness Measures Phase)* | 0.000 | 0.002 | 0.000 |
| State Induction Assessment (Consciousness Measures Type) | 0.000 | 0.002 | 0.000 |
| Binocular Rivalry Task (Task)* | 0.000 | 0.001 | 0.000 |
| Condition Assessment (Consciousness Measures Type)* | 0.000 | 0.001 | 0.000 |
| None (Consciousness Measures Type)* | 0.000 | 0.001 | 0.000 |
| Oddball (Specific Paradigm) | 0.000 | 0.001 | 0.000 |
| None (Stimuli Categories)* | -0.001 | 0.002 | 0.000 |
| Attentional Blink (Specific Paradigm) | 0.000 | 0.001 | 0.000 |
| Sounds (Stimuli Categories)* | 0.000 | 0.001 | 0.000 |
| Sleep Monitoring (Consciousness Measures Type)* | 0.000 | 0.001 | 0.000 |
| PHI (Dependent Measures) | 0.000 | 0.001 | 0.000 |
| Forward Masking (Specific Paradigm) | 0.000 | 0.001 | 0.000 |
| None (Consciousness Measures Phase)* | 0.000 | 0.001 | 0.000 |
| Sleep (Main Paradigm)* | 0.000 | 0.001 | 0.000 |
| Letters (Stimuli Categories) | 0.000 | 0.001 | 0.000 |
| Passive Exposure (Main Paradigm) | 0.000 | 0.001 | 0.000 |
| Unresponsive Wakefulness Syndrome (Specific Paradigm)* | 0.000 | 0.001 | 0.000 |
| Tactile (Stimuli Modality) | 0.000 | 0.000 | 0.000 |
| Electric Stimulation (Stimuli Categories)* | 0.000 | 0.000 | 0.000 |

*Table S6.* Importance factors calculated for each parameter used to train the random forest classifier to classify theory-support according to all of the methodological parameters extracted from the experiments in the database, without the exclusion of parameters due to issues related to multicollinearity (see Figure S2a-b for the classification results). Importance values indicate hamming loss importance scores calculated using a permutation method. Features marked with asterisk were removed from the full model for our main analysis due to multicollinearity issues.

**Supplementary Table S7: Feature importance – content consciousness, neuroscientific techniques**

| Feature | 5% | 95% | Mean Importance |
|---|---|---|---|
| TMS (Neuroscientific Techniques) | 0.009 | 0.034 | 0.021 |
| MEG (Neuroscientific Techniques) | 0.006 | 0.019 | 0.012 |
| fMRI (Neuroscientific Techniques) | 0.000 | 0.027 | 0.012 |
| Random Variable | 0.001 | 0.026 | 0.011 |
| EEG (Neuroscientific Techniques) | -0.001 | 0.026 | 0.009 |
| Intracranial EEG (Neuroscientific Techniques) | 0.000 | 0.011 | 0.005 |

*Table S7.* Importance factors calculated for each parameter used to train the random forest classifier to classify theory-support according to the neuroscientific techniques used in experiments focusing on content consciousness. Importance values indicate hamming loss importance scores calculated using a permutation method.

**Supplementary Table S8: Feature importance – content consciousness, experimental paradigms**

| Feature | 5% | 95% | Mean Importance |
|---|---|---|---|
| Random Variable | 0.016 | 0.048 | 0.032 |
| Direct Stimulation (Main Paradigm) | 0.019 | 0.043 | 0.031 |
| Stimulus Degradation (Main Paradigm) | 0.014 | 0.039 | 0.025 |
| Masking (Main Paradigm) | 0.013 | 0.038 | 0.025 |
| Attentional Manipulation (Main Paradigm) | 0.013 | 0.034 | 0.023 |
| Abnormal Contents of Consciousness (Main Paradigm) | 0.014 | 0.032 | 0.023 |
| Expectation (Main Paradigm) | 0.010 | 0.023 | 0.017 |
| Illusions (Main Paradigm) | 0.007 | 0.021 | 0.014 |
| Competition (Main Paradigm) | 0.007 | 0.020 | 0.013 |
| Emotion (Main Paradigm) | 0.006 | 0.020 | 0.012 |
| Resting State (Main Paradigm) | 0.007 | 0.019 | 0.012 |

*Table S8.* Importance factors calculated for each parameter used to train the random forest classifier, to classify theory-support according to the experimental paradigms used in experiments focusing on content consciousness. Importance values indicate hamming loss importance scores calculated using a permutation method.

### Supplementary boxes

**Supplementary Box A: Description of the core principles and main predictions of the four leading theories of consciousness:**

According to the Global Neuronal Workspace (GNW) theory, conscious processing takes place when information is being shared globally in the brain [2–4]. The transition between unconscious and conscious processing is held to occur when a signal processed by specialized neural modules is broadcasted by a central global workspace of interconnected neurons in fronto-parietal and anterior temporal areas [2,4,5]. The broadcasted signal is then amplified, sustained through time, and shared with other specialized modules in a transient non-linear process called 'ignition' [4,6,7]. The mechanism for ignition is dependent on widespread bi-directional connections between the areas that form the workspace and peripheral modules. This connectivity pattern enables recurrent processing loops between distant brain areas and the resulting mobilization and maintenance of information making it accessible for consciousness [7–9]. GNW accordingly predicts global decodability of subjectively experienced contents from frontoparietal areas, during relatively late time windows, and highlights long-distance connectivity between posterior and anterior areas as the hallmark of consciousness [6,9–11].

In contrast, Higher Order Thought (HOT) stands for a family of theories that share the hypothesis that the first-order representation of a stimulus is insufficient for it to be consciously experienced [12–17]. These theories suggest that for one to be aware of such a first-order representation, a further higher-order representation should be activated [17–19]. These theories typically hold that this higher-order representation is mediated by areas in the prefrontal cortex, mostly the dorsolateral and orbitofrontal areas, that re-represent, or index, the first-order representation [12,14–16]. Thus, these areas are claimed to constitute the NCC. Consequently, HOT theories argue that subjective experience, or “what it is like for me” (Brown, 2015) is represented in the prefrontal cortex, while the information itself is represented in the first-order sensory areas. And it is the latter activity – rather than the higher-order representation - that affects one’s behavioral judgments [14,20,21]. Consequently, these theories remain neutral about whether consciousness has any functional significance, calling for experimental designs in which behavioral performance is matched between conscious and unconscious states, so that no behavioral difference could account for the neural differences between these states [22,23].

HOT theories are typically contrasted with first-order theories [24–26], which negate the claim that an additional stage is needed for consciousness, and argue that activity in first-order, sensory areas, is enough for conscious experience. One such theory is Recurrent Processing Theory (RPT), which – as its name reveals - holds recurrent processing to be the key mechanism subserving conscious experience [25,27,28]. According to RPT, feedforward neuronal processes unconsciously encode perceptual features, while horizontal connections and recurrent loops between lower and higher-level brain areas allow consciousness to occur [28]. Such recurrent loops enable the integration and organization of the isolated encoded features into a conscious unified percept [25,26,29]. Thus, like GNW, RPT requires feedback to sensory modules for consciousness to occur. Yet unlike GNW, RPT holds that this feedback rests on localized recurrent processing loops that are claimed to subserve the experienced content [30–32]. The long-range projections from higher level areas are held by RPT to underlie post-perceptual processes, typically task-related ones [29,32]. In addition, RPT hypothesizes that conscious perception depends on plasticity and not only on recurrent processing per se. Arguably, only recurrent processes that change the brain's connectivity patterns can lead to subjective experience. These changes are hypothesized to depend on feedback-based activation of NMDA receptors in low level perceptual areas [25,30,33].

Finally, Integrated Information Theory (IIT) puts forward a theoretical and mathematical account for consciousness, which is grounded in introspection about the phenomenological properties of consciousness [34–36]. The theory is based on five phenomenological axioms about consciousness from which it derives postulates about the physical substrates of consciousness. IIT considers a system to be conscious if and only if it creates integrated information from the intrinsic perspective of the system [34,37,38]. Accordingly, the dynamic internal structure of a system, in terms of causal interactions between its sub-parts, is crucial for the formation of both the content and the level of consciousness the system has at every given moment in time [35,39]. The theory mathematically formulates the amount of integrated information as Φ, claimed to represent the amount or level of consciousness of the system; it further claims that the content of consciousness is identical to a multidimensional space that is the unfolded cause-effect structure of the system [36,40]. Based on auxiliary hypotheses and neural anatomy, IIT predicts that the NCC (of PSC – Physical Substate of Consciousness; Tononi et al., 2016) should reside in posterior parietal areas, commonly referred to as the posterior-hot-zone. This is because the intrinsic connectivity patterns within and between these areas is hypothesized to have the highest capacity for information integration [39,41].

**Supplementary Box B: Geographical division with respect to the theories**

The database we created also allows one to also ask 'meta-scientific' questions about the sociology of the field. For example, below is a world map of the geographical distribution of papers supporting each of the four theories. Two interesting trends regarding the spread of research in the field can be observed; first, most consciousness research is still centered in Europe (see inset of Supplementary Figure S6), followed by the USA, though studies originate also from other countries, in South America and Asia. Second, the prominence of the theories seems to be relatively stable across countries, with GNW being the most supported in 65% (out of 40) of the countries included in this database, and IIT and RPT being the prominent theories in 20% and 15% respectively (no countries were mostly supportive of HOT).

**Supplementary Figure S6: Geographic distribution of papers by authors**

*Figure S6.* Distribution of the papers in the database according to nations extracted from the authors affiliations. The radius of each concentric circle describes the amount of papers supporting each theory.
GNW= red, RPT=green, IIT=yellow, HOT= blue

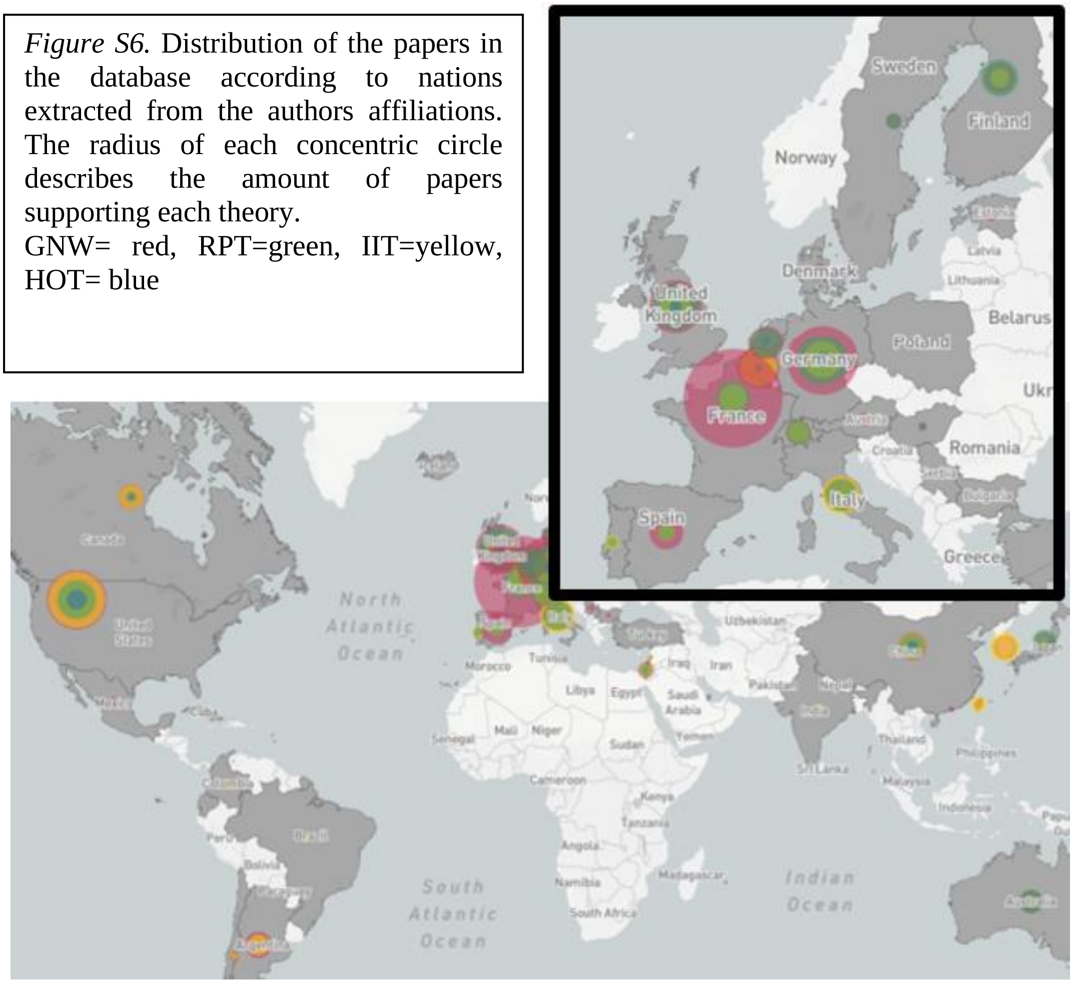
